# A 3D printed model of human lactation

**DOI:** 10.64898/2026.01.30.702762

**Authors:** Amelia Hasenauer, Marcy Zenobi-Wong

**Affiliations:** Tissue Engineering + Biofabrication Laboratory, Department of Health Sciences & Technology, ETH Zürich, Otto-Stern-Weg 7, 8093 Zürich, Switzerland

**Author notes:** **Corresponding author:** Marcy Zenobi-Wong, **Email:**. **Funding:** ETH Foundation Grant 23-1 ETH-12 (M.Z.W.). **Competing interests:** Authors declare that they have no competing interests.

**Keywords:** milk MEC, light printing, microwells, architecture, human lactation

## Abstract

Engineering physiologically relevant breast *in vitro* models remains challenging due to the gland’s complex three-dimensional microanatomy, together with the need for epithelial polarity and hormone responsiveness. To overcome these challenges, fabrication methods are needed that rapidly create alveoli-scale structures with efficient diffusion and sustained hormonal stimulation. Here, Filamented Light (FLight) biofabrication is leveraged to print highly porous, ECM-based hydrogel scaffolds directly within standard Transwell inserts with separate apical and basal access. FLight’s speckle-patterned laser generates multiscale scaffold architectures that integrate filament-derived microchannels (∼15 μm) to promote diffusion with alveoli-inspired cylindrical microwell arrays (Ø100, Ø150, Ø200 μm) that impose geometric constraints to guide epithelial organization. Each insert is printed in <10 s and incorporates slow-release prolactin microcrystals to provide lactogenic stimulation in situ. Primary human milk-derived mammary epithelial cells (milk MECs) were seeded onto the constructs. There, milk MECs line the printed microwells, establish zona occludens-1-positive tight junctions, and express lactation-associated markers (prolactin receptor and β-casein), alongside milk fat globules and intracellular lipid droplets. Collectively, this rapidly reconfigurable FLight platform enables high-throughput generation of hormone-responsive human mammary microtissues for lactation-focused studies and is adaptable to other lumen-forming epithelia.

## 1. Introduction

Lactation is a highly coordinated physiological process that arises from the interplay between maternal endocrine cues and the specialized architecture of the mammary gland.[1] In the breast, clusters of alveoli embedded within the surrounding stroma form the functional secretory units, each organized as a bilayered epithelium with basal and luminal layers.[2,3] Contractile myoepithelial (basal) cells form an outer layer that provides polarity cues and the mechanical force required for milk ejection. Luminal epithelial cells are positioned apically and secrete milk components into the alveolar lumen.[4] This organization enables directional transport across the mammary epithelium. Maternal precursors (e.g., glucose, amino acids, fatty acids, and ions) are taken up basolaterally, converted into milk components (lactose, lipids, and proteins), and secreted apically into the lumen, where milk accumulates and flows through ducts to the nipple.[5] Systemic endocrine cues including prolactin (PRL), oxytocin, and steroid hormones, together with local extracellular matrix (ECM) signals, coordinate epithelial differentiation, milk synthesis, and ejection, to ultimately deliver nutrients, hormones, immunoglobulins, and cytokines tailored to support infant growth and development.[6–8]

The mammary gland’s function emerges from its hierarchical tissue organization that spans the ECM’s ultrastructure (fibril network architecture and ligand presentation) to apical-basal polarity at the level of individual cells and the formation of polarized epithelial layers. This hierarchy further extends to organ-level patterning of spherical lumina (the epithelial hollow spaces that collect secreted milk) and stroma (the supportive connective tissue compartment that surrounds the epithelium and contains fibroblasts, adipocytes, vasculature, immune cells, and ECM).[9] Classic 3D culture studies (such as organoids) in laminin-rich matrices have shown that mammary epithelial cells form β-casein-expressing acini when appropriate apical-basal polarity and basement membrane contact are established.[10–12] By contrast, the same cells in 2D cell culture or disorganized matrices can remain poorly differentiated. Together, these matrix- and geometry-dependent effects show that epithelial behavior is tightly governed by spatial context.[13]

To interrogate how architectural cues or confinement regulate epithelial morphology and function, microwell-based culture systems have been developed. Lithographically defined or molded cylindrical, concave or spherical wells confine small numbers of cells in each microcavity, to promote self-organization into polarized organoids with defined size and composition.[14,15] In hydrogel-based formats, microwell arrays have also been used to generate epithelial layers that line crypt-like wells in intestinal models, to create open, lumen-like compartments with directly accessible apical surfaces.[16] Such configurations are particularly relevant for secretory epithelia such as the mammary gland, where apical access is required to monitor and collect milk secretions.[17] While microwells are physically open at the top, breast epithelial cells cultured in these systems often self-assemble into closed acini with enclosed lumens, and the ability to engineer polarized, apically accessible open lumens with defined architectures remains largely untapped in the context of human lactation models.[14,18]

Current microwell platforms are constrained by how they are fabricated. Many rely on cleanroom-intensive lithography, replica molding, or specialized micromachining, which require dedicated infrastructure and can take hours to days to produce a single batch of devices.[19] These approaches are costly and difficult to reconfigure when new geometries or dimensions are needed.[20] Batch-to-batch variability from multiple molding and surface-treatment steps can further alter well dimensions, wettability, and matrix coatings.[21] Additionally, microwells are typically fabricated separately from the biomimetic ECM. Geometry is first defined in polydimethylsiloxane (PDMS) or thermoplastic masters, and biomimetic ECM cues are introduced only afterward via surface protein coating or secondary hydrogel casting into pre-formed wells, which leads to multi-step assembly workflows. This complicates coordinated control over porosity, stiffness, and biochemical composition and hampers seamless integration into standard labware.[19] Together, these limitations underscore the need for fabrication strategies that can directly generate microwell architectures within biomimetic materials in a single, adaptable process that reduces handling steps and lowers the barrier to broad adoption.

Light-based bioprinting addresses these limitations and enables rapid (seconds-scale) fabrication of biomimetic 3D environments with micrometer-scale control of photo-crosslinking in bioresins (**Figure 1**). Local regions (voxels) in which the accumulated photon dose exceeds a gelation threshold solidify, while unexposed resin is washed away, which yields patterned hydrogels.[17,22] Because printed shapes are defined digitally and projected in parallel, these systems are intrinsically maskless, easily reparametrized for new geometries, and compatible with high-throughput and standard laboratory workflows.[17]

**Figure 1:**
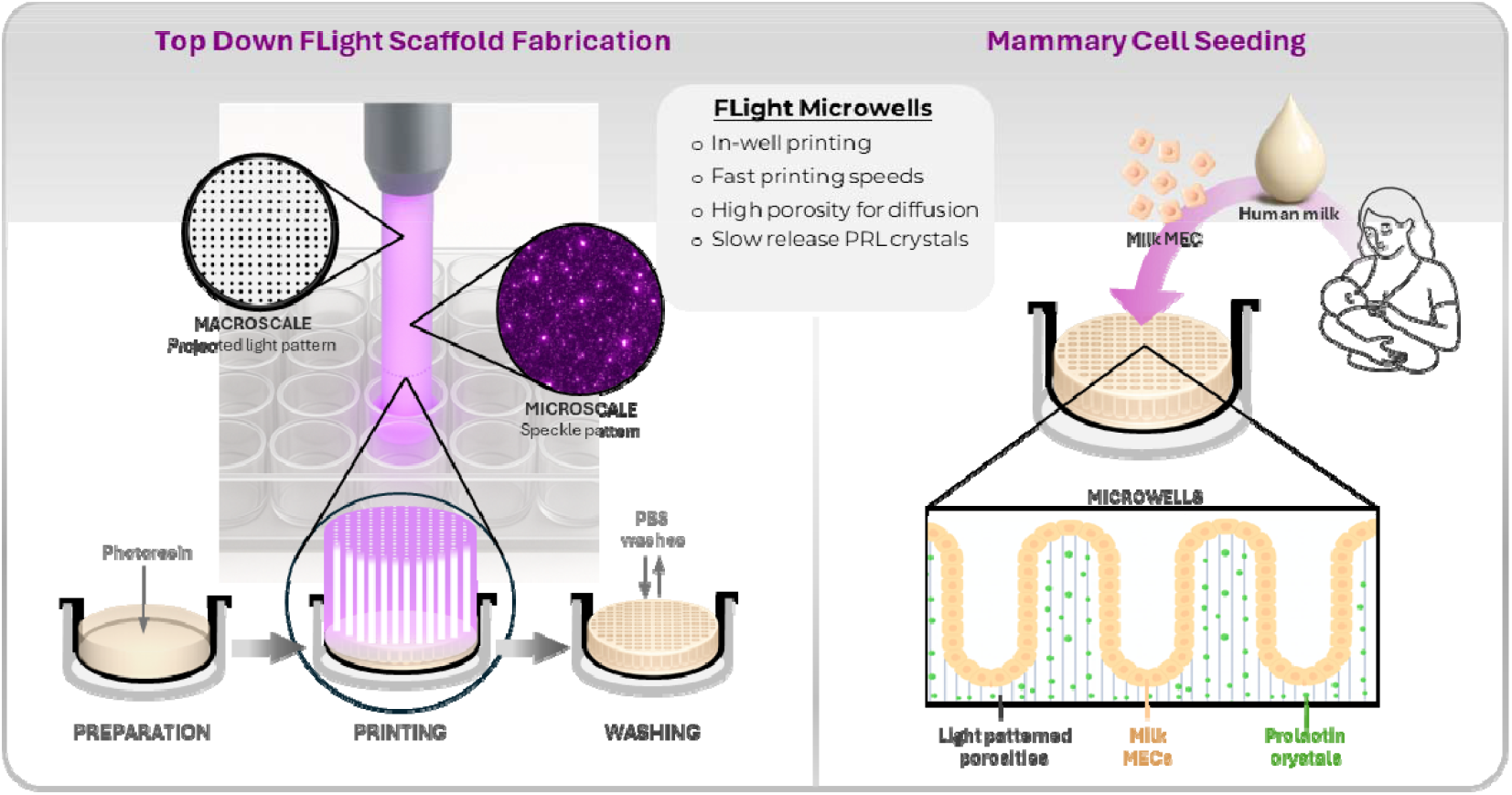
Filamented light (FLight) fabrication of mammary tissue scaffolds. **Left**: A top-down laser projection setup directs speckle-patterned light into standard Transwell inserts, where mammary ECM-composite photoresins are polymerized to form microwell-like architectures with microchannels for diffusion. **Right**: Human milk-derived mammary epithelial cells (milk MECs) are seeded onto the printed microwell scaffolds with prolactin (PRL)-releasing microcrystals to provide lactogenic cues.

Among light-based biofabrication technologies, filamented light (FLight) represents a distinct modality for three-dimensional structuring (**Figure 1**). In FLight, speckle-patterned light is concentrated into self-focused filaments that locally induce polymerization. This yields hydrogel scaffolds with interconnected, high aspect-ratio microchannels (∼2–20 μm in diameter) that can guide cell growth and migration.[23] To date, FLight has been applied primarily to the fabrication of aligned architectures for anisotropic tissue engineering, with emphasis on microchannel-scale control of cellular organization (e.g. muscle, cartilage) and nuclear confinement (e.g. tendon).[24–26] However, FLight can also be extended to pattern macroscale construct geometries with high lateral resolution (∼50–100 μm) and geometric complexity (**Figure 1**).[25,26]

Here, we leverage FLight’s multiscale patterning capability to fabricate ECM-based, alveoli-scale architectures (macroscale) with speckle-derived microchannels (microscale) to enhance nutrient transport. This integrated architecture supports the formation of a lactation-relevant mammary epithelial tissue model. Using top-down laser projection, FLight printed scaffolds are formed directly within standard Transwell inserts, with distinct apical and basal compartments that capture key architectural features of the lactating breast and provide a generalizable framework for modeling other secretory epithelia.

## 2. Results

### 2.1. ECM-composite photoresins for light-based mammary scaffold printing

Collagen I (Col-I), Matrigel, and mammary-derived decellularized extracellular matrix (dECM_mam_) represent complementary ECM sources for engineering mammary tissue-mimetic hydrogels. Col-I is a major structural component of the mammary stroma and provides a fibrillar backbone, while Matrigel supplies basement membrane-associated proteins. Both are commercially available and widely used. In contrast, dECM_mam_ provides organ-specific ECM complexity (e.g. laminins, glycoproteins and proteoglycans) and supplies tissue-relevant biochemical cues that are difficult to reproduce with purified components alone. [17]

Thus, photoresins for FLight printing of mammary scaffolds were prepared in three formulations: (i) Matrigel/Col-I (5 mg/mL Matrigel + 5 mg/mL collagen I), (ii) dECM_mam_/Col-I (20 mg/mL bovine dECM_mam_ + 5 mg/mL Col-I), and (iii) Matrigel/dECM_mam_ (5 mg/mL Matrigel + 20 mg/mL bovine dECM_mam_) (**Figure 2A**). All formulations contained the photoinitiator pair ruthenium/sodium persulfate (Ru/SPS; 0.1 mM/10 mM) to enable visible-light-induced oxidative crosslinking of tyrosine residues and formation of stable polymer networks (**Figure 2A**).

**Figure 2:**
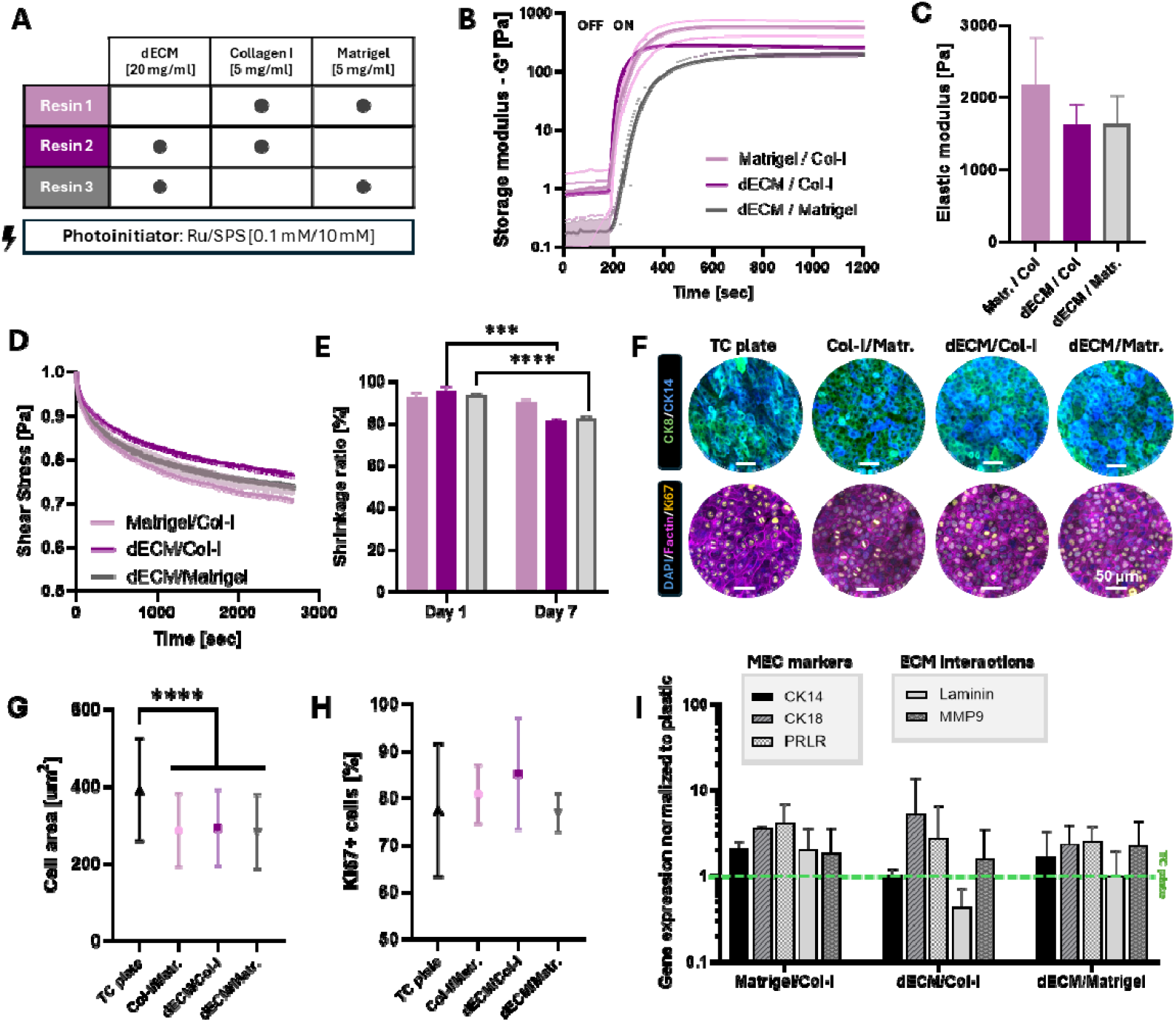
Characterization of photoresins for FLight mammary tissue engineering. **A)** Composition of the three photoresin formulations: bovine udder decellularized extracellular matrix (dECM_mam_), collagen I, and/or Matrigel with Ru/SPS photoinitiator. **B)** Light induced gelation of the three photoresins shown by photorheology (405 nm light). **C)** Elastic modulus of fully crosslinked hydrogels. **D)** Stress-relaxation profiles show viscoelastic behavior and faster stress dissipation in dECM-containing hydrogels. (Mean ± S.D, n = 3 technical replicates for panels B-D) **E)** Shrinkage over 7 days, showed minimal deformation for all three formulations; Resin 1 - violet, Resin 2 - purple, Resin 3 - grey (Mean ± S.D, n = 5 technical replicates, p<0.001^***^, p<0.0001^****^). **F)** Confocal images of MCF10A cells cultured on standard tissue culture plates (TC plates) or on hydrogels for 4 days, stained for CK8 (green) and CK14 (blue) (top), and F-actin (pink), Ki67(yellow) and DAPI (blue) (bottom). Scale bars: 50 μm. **G)** MCF10A cell area quantification on mammary hydrogels compared to TC plastic controls. **H)** Percentage of Ki67□proliferating cells. **I)** Gene expression of differentiation and ECM-interaction markers in MCF10A cells on photoresins, normalized to TC plastic (green dashed line). Mean ± S.D, n = 3 independent cell cultures for panels G-I, p<0.0001^****^.

Photorheological measurements under a 405 nm illumination demonstrated rapid, light-induced gelation across all three formulations (and their single components), with plateau storage moduli (G′) of □600 Pa for Matrigel/Col-I□300 Pa for dECM_mam_/Col-I, and □180 Pa for Matrigel-dECM_mam_ (**Figure 2B** and **S1A, B**, Supplementary Information). Matrigel/Col-I hydrogels exhibited the highest elastic modulus (∼2 kPa), whereas dECM_mam_-containing formulations were softer (∼ 1.5 kPa), although no statistical significant differences were observed (**Figure 2C**). All hydrogels were viscoelastic and stress relaxation occurred more rapidly in dECM_mam_-based formulations compared to the Matrigel/Col-I composite (**Figure 2D**). Over a seven-day incubation at 37 °C in PBS, the three crosslinked mammary hydrogels were stable and showed only limited shrinkage, with approximately 20% contraction in dECM_mam_-based hydrogels and 10% in Matrigel/Col-I composites (**Figure 2E** and **S1C**, Supplementary Information).

To test whether the matrices support mammary epithelial cell (MEC) adhesion and epithelial phenotype, non-tumorigenic human breast epithelial MCF10A cells were cultured on the crosslinked hydrogels and on standard tissue culture (TC) plastic as controls. Across all hydrogel formulations, cells attached and proliferated to form epithelial monolayers expressing cytokeratin 8 (CK8, luminal marker) and cytokeratin 14 (CK14, basal marker) (**Figure 1F**). Quantitative analysis showed that MCF10A cells cultured on tissue culture plastic had a significantly larger spread area than cells cultured on the hydrogel matrices (**Figure 1F, G**). In contrast, cells on hydrogels displayed a more compact morphology and formed denser cell clusters. This trend is consistent with prior reports showing that compliant matrices and matrix-specific cell–ECM interactions limit adhesion driven epithelial cell spreading compared with rigid 2D plastic surfaces.[27] Proliferation of MCF10A cells, assessed by Ki67 expression, was comparable across all conditions and TC plastic (**Figure 2F, H**). At the transcriptional level, the photoresins promoted the expression of key mammary epithelial genes. RT–qPCR revealed higher transcript levels of cytokeratins (CK14 and CK18) and the prolactin receptor (PRLR) in cells cultured on the hydrogels compared with tissue-culture plastic (**Figure 2I**). Expression of ECM interaction-associated transcripts, with matrix metalloprotease 9 (*MMP9*) and *LAMININ*, was generally above the TC plastic baseline for all mammary hydrogels. The only exception was *LAMININ* in the dECM_mam_/Col-I condition which remained below the tissue-culture plastic baseline (**Figure 2I**).

These data establish the mammary ECM-composite resins as robust, rapidly photo-crosslinkable matrices that support MEC phenotype and gene programs, for light-based printing of mammary scaffolds.

### 2.2. FLight printing of aligned and porous hydrogel microstructures improves molecule diffusion

FLight printing was performed with a custom-built top-down projection setup. A 405 nm speckled laser was guided through an aperture and optics to be projected onto a digital micromirror device (DMD), and redirected “top-down” by a mirror into glass cuvettes or well plates to crosslink the mammary ECM-composite resins in user defined patterns (**Figure 3A**).

**Figure 3:**
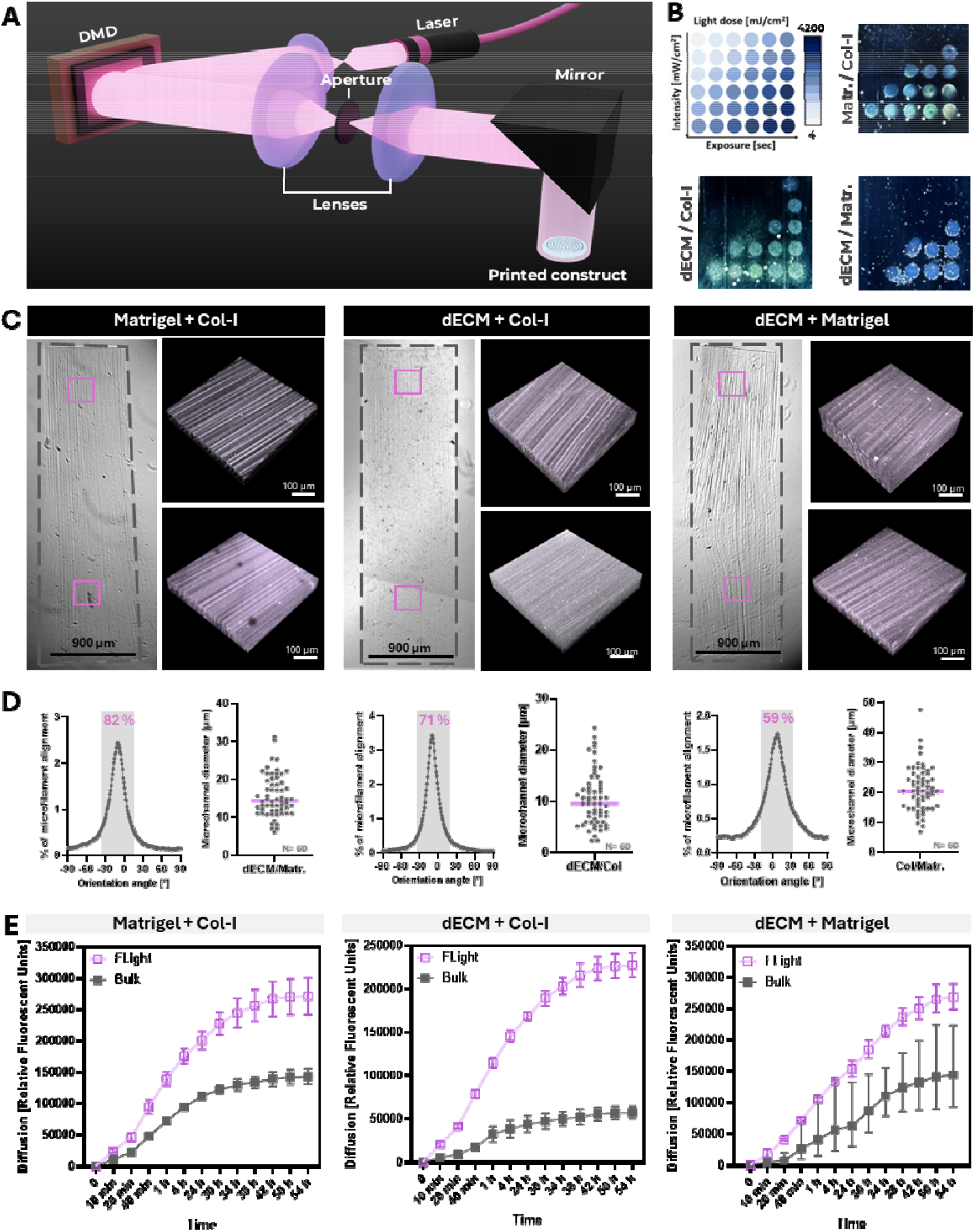
FLight printing generates aligned, porous micro-scale architectures in mammary ECM-composite photoresins. **A)** Schematic of the custom top-down FLight printer: Laser (405 nm) projection patterns were modulated with a digital micromirror device (DMD) and relayed through a lens-aperture assembly and mirror optics to project spatially patterned structures onto the mammary resin during fabrication. **B)** Light-dose matrices (intensity × exposure time) projected onto Matrigel/Col-I, dECM_mam_/Col-I and dECM_mam_/Matrigel photoresins. Each dot (1 × 1 mm) represents one tested light dose for printing. Exposure time range: 4-64 sec; Light intensity range: 2-64 mW cm^-2^. **C)** Bright-field images and confocal reconstructions of millimeter-scale cylinders printed from each formulation. Constructs contained dense and continuously aligned filament networks (dashed boxes delimit the cylinder) Scale bars: 900 μm (left) and 100 μm (right). **D)** Histograms of filament angle orientation (alignment), with the percentage of filaments within ±15° of the dominant axis (**left**); quantification of filament-to-filament spacing (void spaces) for each resin (n = 60 void spaces from three technical replicates). Scale bars: 50 μm (**right**). **E)** Diffusion kinetics (10 kDA Rhodamine) in the three mammary hydrogels crosslinked with FLight printing or with homogenous UV light (bulk) at matched light intensities.

FLight printing was optimized with a matrix of light intensities and exposure times to identify an optimal printing light dose for each photoresin (**Figure 3B** and **S2A**, Supplementary Information). The final formulation-specific doses were 625.5 mJ cm^−2^ for dECM_mam_/Matrigel, 334.6 mJ cm^−2^ for dECM_mam_/Col-I, and 125.5 mJ cm^−2^ for Matrigel/Col-I.

Millimeter scale pillars were printed, which exhibited FLight’s characteristic striations (microfilaments).[23] This anisotropic structure spanned the full cylinder width and length (**Figure 3C**). Orientation maps confirmed robust alignment of microfilaments across all formulations, with the highest alignment in Matrigel/Col-I (82%), followed by dECM_mam_/Col-I (71%) and dECM_mam_/Matrigel (59%) (**Figure 3D**, left and **S2B**, Supplementary Information). A technical consideration is that residual dECM particles may modestly interfere with filament-alignment quantification. As previously shown, microfilaments generated by FLight patterning were separated by aligned microchannels oriented along the printing direction.[23] Microchannel diameter measurements in mammary hydrogels were in the tens-of-micrometers range and varied by formulation: dECM_mam_/Col-I, 10.5 ± 5.3 μm (range 2.4–24.3 μm); dECM_mam_/Matrigel, 15.6 ± 5.5 μm (6.1–31.2 μm); Matrigel/Col-I, 21.1 ± 7.4 μm (6.8–47.6 μm) (**Figure 3D**, right). To test whether these aligned porosities improved molecular transport, diffusion was quantified in FLight versus bulk hydrogels. FLight-printed constructs showed higher diffusion than their bulk controls for each formulation, with the strongest divergence emerging after 1 h (**Figure 3E**). dECM_mam_/Matrigel hydrogels, which combined larger microchannel diameters with lower stiffness, exhibited the highest cumulative diffusion levels. Overall, these findings indicate that diffusion in FLight-patterned mammary hydrogels reflects both bulk matrix properties and filament-induced microarchitecture, where aligned microchannels contributed low-resistance transport pathways.

### 2.3. Alveoli-scale macrostructure FLight-printing and feature fidelity

Across mammals, the secretory parenchyma of the pregnant or lactating mammary gland is organized into densely packed lobuloalveolar units composed of spherical alveoli.[28] To define physiologically relevant dimensions for printed mammary architectures, alveolar diameters were assessed in histological sections of human breast and lactating bovine udder tissue (**Figure 4A**, top). Human breast alveoli measured 56.9 ± 16.3 μm (median 54.8 μm), whereas lactating bovine udder alveoli measured 64.3 ± 20.5 μm (median 61.3 μm). Lactating bovine alveoli had a significantly larger mean diameter than human breast alveoli (_Δmean_ = 7.4 μm), which reflected potential differences in lactation stage or species-specific gland architecture.[28] Considering both tissues, native alveolar diameters spanned ∼40–120 μm, which provided a physiologic target range for alveoli-scale feature resolution in the printed constructs (**Figure 4A**, bottom).

**Figure 4:**
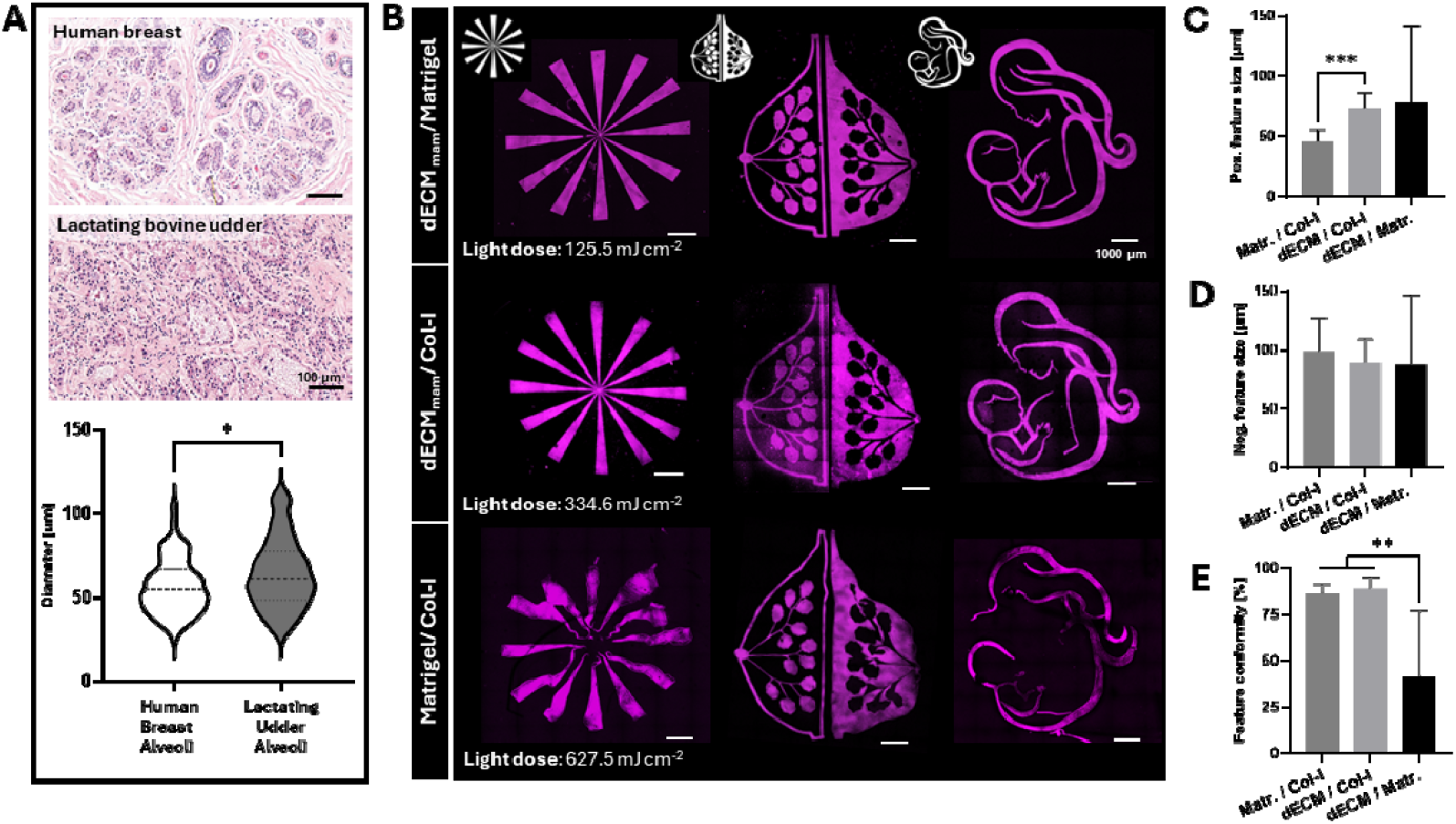
Top-down FLight printing of mammary ECM-composite resins at high macroscale resolution. **A)** Histological sections of human breast and lactating bovine udder tissue (Hematoxylin & Eosin stained) and quantification of native alveolar diameters (n = 100 alveoli, p^*^<0.05, Scale bar: 100 μm). **B)** Representative Top-Down FLight prints of a spokes wheel and complex mammary-inspired geometries fabricated in three ECM-composite photoresins composed of 20 mg/ml dECM_mam_, 5 mg/ml Matrigel and 5mg/ml Col-I printed with formulation-specific light doses of 627.5, 334.6, and 125.5 mJ cm^−2^. Scale bar: 1000 μm. **C)** Quantification of positive and **D)** negative feature sizes for each formulation. **E)** Feature conformity as percent agreement between designed and printed features (Mean ± S.D, n = 9 with 3 prints measured 3 times. ^**^p<0.01, p<0.001^***^).

Top-down FLight projection printing was assessed for its ability to generate *in vivo*–scale features across the three ECM-composite photoresins. A spoked-wheel resolution pattern and complex mammary-inspired geometries were printed using dECM_mam_/Matrigel, dECM_mam_/Col-I, and Matrigel/Col-I photoresins at formulation-specific light doses of 627.5, 334.6, and 125.5 mJ cm^−2^, respectively (**Figure 4B**). All constructs were subjected to the same PBS washing workflow prior to imaging and quantification. The resulting prints indicated sharper edges and improved feature continuity for the collagen-containing formulations, whereas dECM_mam_/Matr exhibited deformed and discontinuous features post printing and washing (**Figure 4B**). Quantitative analysis confirmed formulation-dependent limits for positive (solid) feature resolution. Matrigel/Col-I produced the smallest positive features (45.7 ± 9.8 μm), whereas dECM_mam_/Col-I and dECM_mam_/Matrigel yielded larger features (73.4 ± 12.3 μm and 77.9 ± 63.9 μm, respectively (**Figure 4C** and **S3A**, Supplementary Information). In contrast, negative (void) feature sizes were comparable across groups with dECM_mam_/Matrigel 87.4 ± 59.2 μm, dECM_mam_/Col-I 89.1 ± 20.1 μm, and Matrigel/Col-I 98.7 ± 28.0 μm (**Figure 4D** and **S3A**, Supplementary Information). Feature conformity after PBS washing differed markedly, where dECM_mam_/Matrigel showed low and variable agreement with the design (41.4 ± 35.6%), and the collagen-containing resins maintained high conformity (dECM_mam_/Col-I, 88.7 ± 6.5%, and Matrigel/Col-I, 86.2 ± 5.0%) (**Figure 4E**). Notably, dECM_mam_/Matrigel required the highest exposure (627.5 mJ cm^−2^) yet still underperformed relative to collagen-containing formulations printed at lower doses (**Figure 4B, E** and **S3B, S4**, Supplementary Information). Accordingly, subsequent scaffold printing and cell studies were performed using Matrigel/Col-I (Resin 1) and dECM_mam_/Col-I (Resin 2).

### 2.4. Geometry-guided mammary epithelial morphogenesis in lumenized scaffolds

To establish geometric control over mammary epithelial cells, cylindrical channel (microwell) arrays were printed with diameters approximating the in vivo alveoli-scale (Ø100, Ø150, Ø200 μm) directly into 24-well Transwell inserts using Matrigel/Col-I and dECM_mam_/Col-I photoresins. After PBS washing, the microwells remained well resolved across all designs as confirmed by confocal imaging (**Figure 5A** and **S5A**, Supplementary Information). The resulting feature sizes closely matched the nominal designs in both photoresins. Matrigel/Col-I yielded 110.9 ± 15.9 (73.5–153.2), 152.9 ± 12.4 (114.7–172.1), and 206.2 ± 17.6 (170.7– 239.0) μm for Ø100, Ø150 and Ø200 μm, respectively, and dECM_mam_/Col-I yielded 111.2 ± 7.7 (96–130), 146.9 ± 14.6 (110–168), and 210.8 ± 13.7 (179–234) μm (**Figure 5A** and **S5B, C** Supplementary Information). Under a ±10% acceptance band (Ø 100: 90–110 μm; Ø150: 135–165 μm; Ø200: 180–220 μm), both resins met the Ø200–150 nominal specifications on average, whereas both resins exceeded the Ø100 specification on average. Channel depth was controlled by the hydrogel fill height within the insert and was set to 500 μm for this study to produce alveolar-like features. Increasing the fill height to 1 mm generated more duct-like architectures (**Figure 5** and **S6**, Supplementary Information).

**Figure 5:**
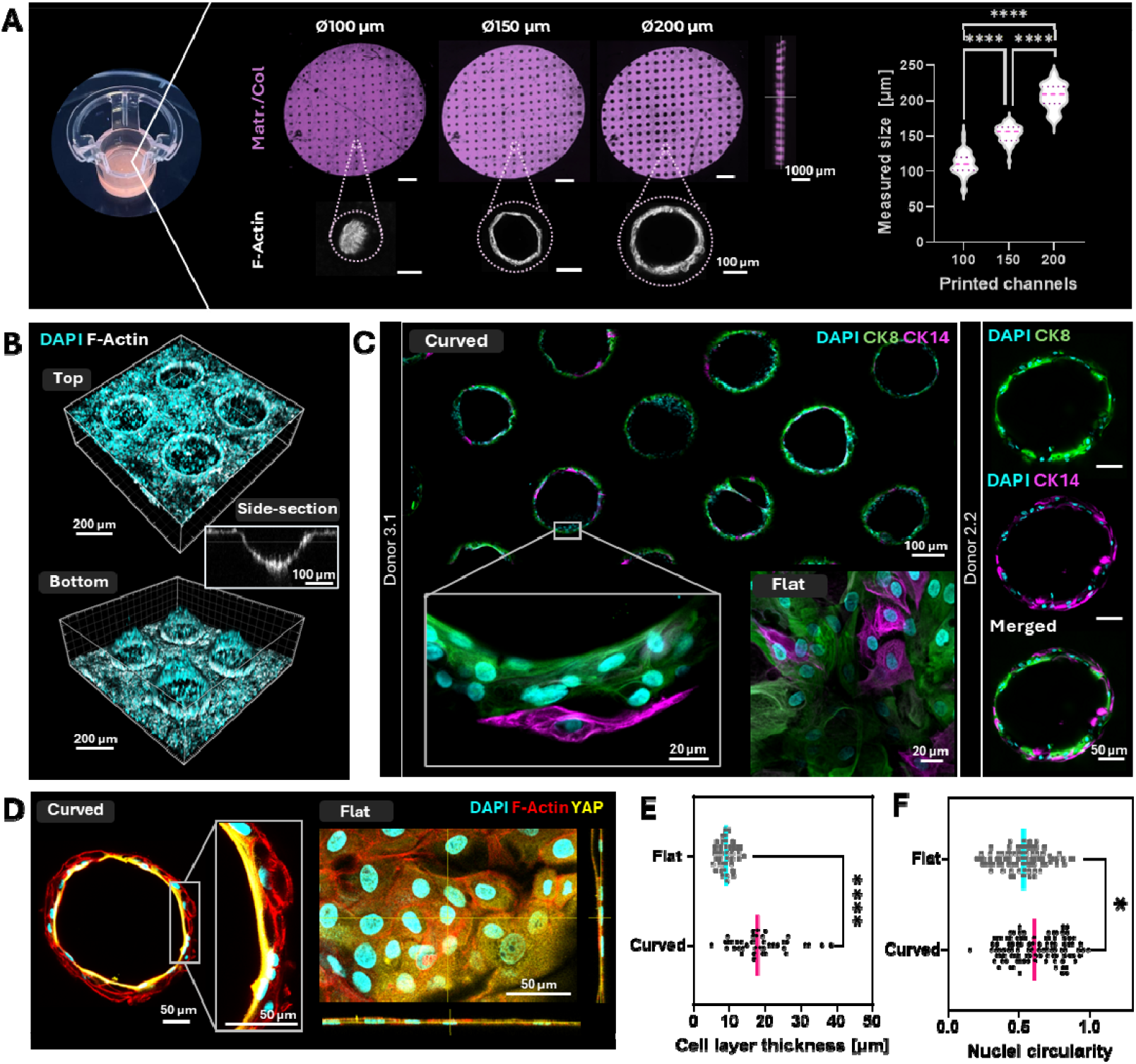
Milk-derived mammary epithelial cells self-organize into epithelial-lined lumens within printed microwell arrays. **A**) left: Picture of a FLight microwell insert printed within a Transwell insert. Middle: representative Matrigel/Col-I (purple) channel arrays (100, 150 and 200 μm nominal diameters), with a representative z-view of Ø200; and representative confocal images of milk MEC (stained with F-actin, grey) cultured on the microwell arrays. Right: Measured channel diameters for each printed design. **B)** 3D reconstruction of the Ø200 μm insert colonized by milk MEC (DAPI: cyan; F-actin, white). **D)** Immunofluorescence imaging of luminal (Cytokeratin 8) and basal (Cytokeratin 14) milk MECs within the channel-lining epithelium (Donor 2.2 and Donor 3.1). **E)** Curved versus flat regions stained for DAPI (cyan), F-actin (red), and YAP (yellow), highlighting curvature-associated differences in epithelial organization and nuclear morphology. **F)** Quantification of nuclear circularity and epithelial layer thickness in flat versus curved regions. (Mean ± S.D, n = 100 measurements from 5 technical replicates, ^*^p<0.05, p<0.0001^****^).

After initial material optimization using MCF10A cells (**Figure 2)** and a preliminary cytocompatibility assessment of stromal cells encapsulated within the matrix (**Figure S7**, Supplementary Information), studies were extended to primary milk-MECs to test the printed geometries in a more lactation-relevant epithelial population (**Table S1**, Supplementary Information).[29,30] Milk MECs isolated from breastfeeding donors were seeded into microwell arrays spanning Ø100 to >Ø200 μm to define the lower geometric limit for reproducible lumen formation *in vitro* (**Figure 5** and **S8A**, Supplementary Information). Milk MECs occluded Ø100 μm channels, while Ø150 μm channels showed variable lumen formation with occasional clogging. Ø200 μm channels more consistently supported open, epithelial-lined lumens and were therefore selected as the standard geometry to maximize robustness across donors and time points for subsequent studies (**Figure 5A** and **S8**, Supplementary Information). Orthogonal slices of the Ø200 μm wells revealed a rounded, bowl-shaped base in the z-dimension rather than a perfectly cylindrical microwell. This geometry likely reflects a combination of depth-dependent polymerization effects (e.g., reduced light penetration) and cell-mediated contractility, consistent with epithelial smoothing of sharp features in response to corner sharpness and curvature-dependent boundary mechanics (**Figure 5B**).[31] No epithelial cell migration into the FLight-generated microchannels was observed. Under the conditions tested, cells remained confined to the printed macroarchitectures and formed continuous epithelial linings (**Figure 3, 5B** and **S9**, Supplementary Information).

Human milk MECs cultured within the printed channels expressed both luminal (CK8) and basal (CK14) cytokeratins along the epithelialized walls, consistent with maintenance of mammary epithelial identity after seeding and culture (**Figure 5D**). Higher-magnification images suggested partial spatial segregation, where selected regions showed CK8^+^ cells at the lumen-facing side and CK14^+^ toward the matrix-facing surface. Overall, organization remained discontinuous and complete spatial self-reorganization into a fully resolved bilayer was not achieved under current conditions (**Figure 5D**).

Flat and curved scaffold regions were compared to assess geometry-dependent epithelial morphology and mechanosensitive signaling. Flat regions corresponded to the planar areas between adjacent microwells, whereas curved regions comprised the concave inner walls of the microwells. F-actin and Yes-associated protein (YAP) staining showed lumen-facing enrichment of YAP in curved segments (**Figure 5E**), consistent with prior reports that tissue geometry can influence YAP subcellular distribution.[16,32] Curved regions exhibited greater epithelial thickness (18.8 ± 7.5 μm) than flat regions (9.1 ± 2.1 μm) (**Figure 5E** and **S10**, Supplementary Information). Nuclear morphology also differed, with higher circularity in curved regions (0.58 ± 0.18) compared with flat regions (0.52 ± 0.16) (**Figure 5F**).

### 2.5. Matrix-embedded PRL-depots support lactogenic activation of milk MEC

To move toward a fully integrated lactation scaffold, endocrine stimulation was next directly integrated within the matrix, rather than relying on lactogenic hormones delivered via the culture medium.[17] Prolactin is a principal lactogenic hormone that promotes mammary epithelial differentiation and milk-protein gene expression through PRLR/Janus kinase 2/Signal Transducer and Activator of Transcription 5 (STAT5) signaling.[33,34]

A “lactation resin” was formulated that contained slow release PRL microcrystals (PODS - Polyhedrin Delivery System) (**Figure 6A** and **S11A**, Supplementary Information). PODS-loaded resins were FLight printed at high resolution, with sharply defined features, which indicated that their inclusion remained compatible with the printing process. The PODS were distributed throughout the printed hydrogel (**Figure 6A, B** and **S11B**, Supplementary Information).

**Figure 6:**
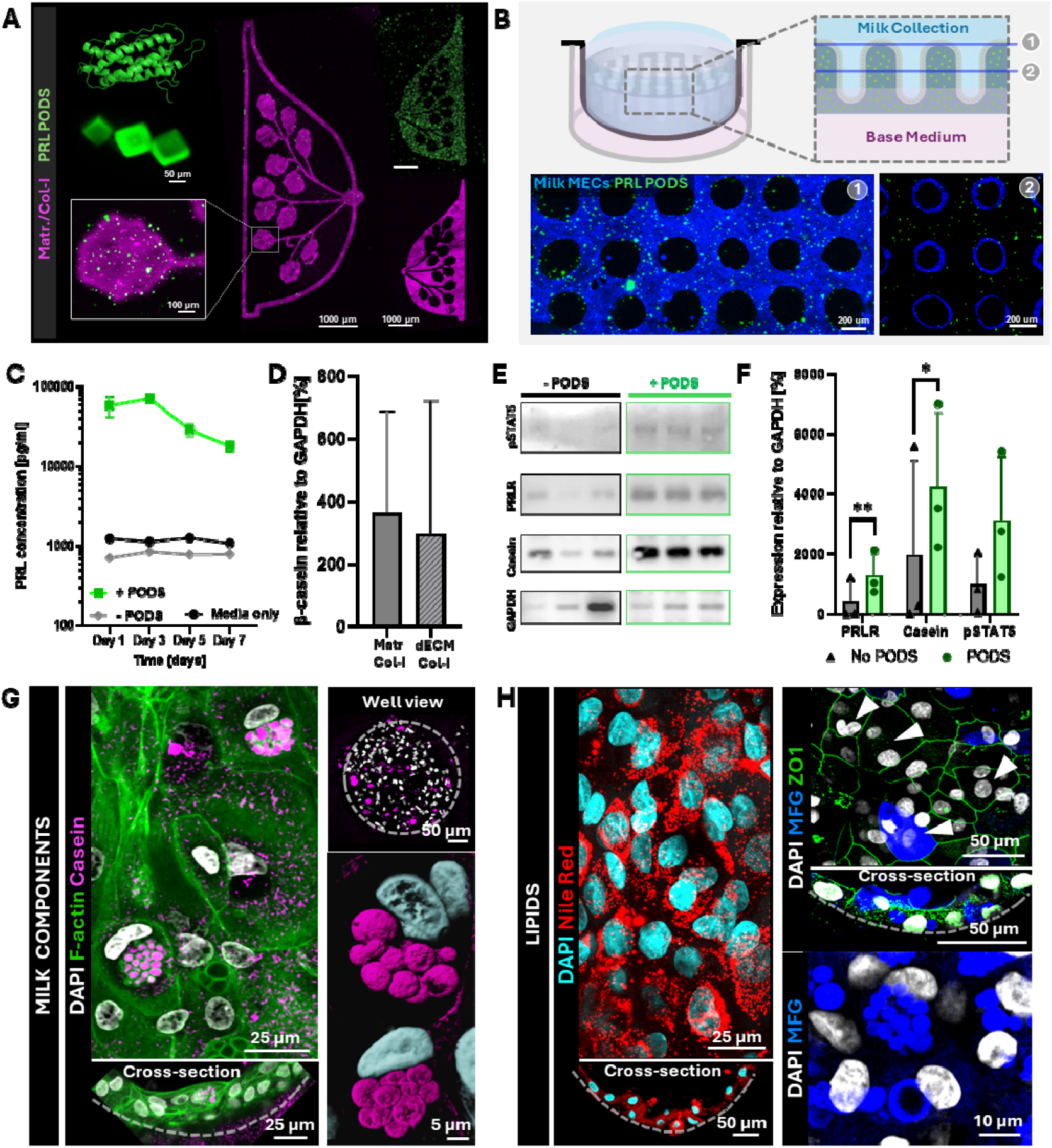
Printed PRL PODS–loaded “lactation resin” enables sustained prolactin release and induces lactogenic signaling and milk-associated outputs in engineered lumina. **A)** PRL (Nuclear magnetic resonance structure*[35]*) and representative images of PRL loaded PODS. PODS (green) were resuspended in ECM-composite photresins (Matrigel/Col-I, purple) to generate “lactation resin”, that were printed at high resolution. **B)** Microwells (Ø200 μm) were printed with the lactation resin into a Transwell to form two-compartment setup (milk-collection chamber above base medium). Representative fluorescence images (top view and cross-section) show milk MECs (Cellmask, blue) and PRL-PODS (green) within Ø200 μm microwell scaffolds. **C)** PRL release profile from PRL-PODS-containing Ø200 μm microwell scaffolds (+ PODS) over 7 days compared with scaffold without PODS (−PODS) and media only controls. **D)** β-casein expression in constructs printed with Matr/Col-I or dECM_mam_/Col-I, quantified relative to GAPDH. **E)** Western blots for phosphorylated STAT5 (pSTAT5), prolactin receptor (PRLR), beta casein, and GAPDH scaffolds printed and cultured without (−PODS) or with (+PODS) PRL-PODS. **F)** Densitometric quantification of PRLR, casein, and pSTAT5 normalized to GAPDH for −PODS and +PODS conditions. **G)** Immunofluorescence staining of milk protein and cytoskeletal organization in lactation Ø200 μm microwell scaffolds (+PODS), showing DAPI (nuclei, white), F-actin (green), and casein (pink), and **H)** Immunofluorescence staining of lipid-associated milk components such as neutral lipids (Nile Red) and milk fat globules (MFG; MFG-E8). Tight junctions localized on the apical side were stained with ZO-1. Mean ± S.D, n = 3, p<0.05 and ^*^p<0.01)

Microwells (Ø200 μm) were printed from the lactation resin directly within Transwell inserts to establish a two-compartment culture configuration. Medium was supplied to the basal chamber in contact with the PODS-containing scaffold, and secreted milk-associated products were collected from the apical chamber (**Figure 6B**). Milk MECs were seeded onto the printed scaffold and the matrix-embedded PRL-PODS released prolactin over 7 days in a degradation-dependent manner and thereby presented the PRL hormone to the basal/basolateral of the epithelial layer (**Figure 6C**). This configuration better reflects physiological endocrine presentation from the stroma and basement membrane than uniform prolactin supplementation in the culture medium.[3] In both Matrigel/Col-I and dECM_mam_/Col-I scaffolds with PODS, the lactation output of milk MEC assessed by beta casein and PRLR expression was comparable, with no statistically significant differences between matrices (**Figure 6D** and **S12**, Supplementary Information).

In milk MEC seeded scaffolds the inclusion of PRL-PODS led to a significantly higher PRLR and β-casein protein expression, together with an increase in phosphorylated STAT5 (pSTAT5), compared with PODS-free scaffold controls (**Figure 6E** and **F**). In MCF10A, PRLR also significantly increased with PODS, while changes in β-casein and pSTAT5 were smaller and did not reach significance under these conditions (**Figure S13**, Supplementary Information). Overall, PRLR, pSTAT5, and β-casein expression was higher in milk MECs than in MCF10A, which supported the use of milk MEC as the preferred model for assays requiring lactation-associated protein expression (**Figure S13**, Supplementary Information), consistent with reports that MCF10A cells are more basal-like and can exhibit atypical differentiation that may not fully reflect normal mammary epithelial function.[36,p.1]

Given the induction of lactation-associated proteins in milk MEC scaffolds, the spatial distribution of lactation associated products was examined. At day 7 on PODS-containing microwells, β-casein was expressed within epithelial regions as droplet-like structures rather than as diffuse cytoplasmic distribution, which suggested localized β-casein accumulation (**Figure 6G**). Neutral lipid staining (Nile Red) showed abundant puncta throughout milk MEC regions with enrichment near the lumen boundary (**Figure 6H**), and milk fat globule (MFG; MFG-E8) signal was detected as both diffuse cytoplasmic staining and rounded droplets (**Figure 6H**). In parallel, milk MECs formed apical tight junctions (ZO-1), consistent with epithelial polarization (**Figure 6H**). Binucleated cells were also observed within the printed scaffolds (**Figure 6H**), a feature reported in lactating mammary epithelium and associated with lactation-linked polyploidization.[17,37]

## 3. Discussion and Conclusion

Here we present a human lactation-focused *in vitro* model which combines a polarized mammary epithelium, apical access for sampling, sustained endocrine stimulation, and biomimetic ECM materials. We adapted top-down filamented light (FLight) printing to generate porous microwell arrays for mammary tissue modeling. Scaffolds were fabricated within seconds (<10 s/scaffold) and integrated into standard workflows by printing directly into commercially available well plates and Transwell inserts, which supported straightforward adoption in routine workflows and higher-throughput experimentation. The resulting scaffolds combined digitally defined, alveoli-scale features with filament-induced micron-scale porosity, which increased molecular diffusion compared with bulk-crosslinked hydrogels.

Milk-derived MEC offer a non-invasive, human cell source for studying mammary epithelial behavior *in vitro*.[17] After isolation from human milk and expansion, cells seeded into Ø200 μm microwell arrays formed open, epithelial-lined lumina and showed curvature-associated changes such as epithelial thickening, increased nuclear circularity, and mechanosensitive signaling (shift in YAP distribution toward the lumen-facing side). Across experiments, Ø200 μm wells were the most reliable for maintaining open lumina, whereas smaller wells more often occluded, which underscored that physiologic feature sizes do not automatically yield reproducible architectures *in vitro*. To align our platform with lactation applications, we embedded slow-release PRL microcrystals (PODS) directly within the photoresin. This yielded a microwell “lactation unit” in which endocrine cues are built into the matrix. Because hormone release occurs from within the scaffold, PRL exposure is expected to be biased toward the basolateral interface, more consistent with endocrine delivery from the stromal side than uniform hormone supplementation in the media. Milk MEC cultured on these platforms exhibited lactation-associated features, such as PRLR, β-casein and pSTAT5 expression, as well as milk-fat–associated readouts (MFG-E8 signal and neutral lipid accumulation) and ZO-1–positive apical junctional organization, consistent with polarized epithelial states.

However, in the reported experiments, milk-associated proteins were detected in both flat and curved regions, which indicated that curvature may not be required for lactation-associated protein expression under the conditions tested. This raises the question of whether alveolar curvature directly influences milk synthesis/secretion or instead reflects a mechanically stable geometry that emerges during epithelial self-organization. Rounded lumina can lower surface-related energy and distribute tension more evenly, and many lumenized tissues (e.g., salivary, pancreatic acini, epithelial cysts, thyroid follicles) adopt near-spherical shapes, which suggests that curvature may often arise from general physical constraints rather than being optimized specifically for lactation.[38–40] Consistent with this interpretation, the curvature-dependent differences we observed in epithelial thickness and YAP localization support the idea that geometry can modulate organization and mechanotransduction in our system, however, these data do not establish that curvature is necessary for milk production. The link between curvature, epithelial organization and secretory output is also relevant for translation toward cell-based milk production. Because planar perfusion and hollow-fiber reactors impose different geometry and transport/collection conditions, they may differentially affect epithelial stability, mass transfer, and product recovery.[41–43] Here our platform could serve as a microscale testbed to probe bioprocess-relevant readouts such as apical product accumulation, sensitivity to transport limits, and lipid/protein retention versus release to help inform design considerations for scale-up.

To test whether curvature has an independent effect on lactation, tighter control over 3D geometry will be needed, such as systematic changes in out-of-plane curvature. Our current top-down, single-projection approach prioritizes speed and is effectively 2.5D, with high lateral fidelity (Ø 200 μm wells printed at 206.2 ± 17.6 μm in Matrigel/Col-I and 210.8 ± 13.7 μm in dECM_mam_/Col-I) but limited depth control, and therefore does not currently enable printing tapered or fully 3D curved lumina. Future improvements could include z-resolved patterning with future hardware developments of the FLight printer allowing sequential exposures, or complementary methods such as molding or two-photon ablation.[44] In addition, denser (honeycomb-like) array layouts could reduce large flat regions that dilute bulk readouts, provided wall stability and feature separation are maintained.[45]

Although milk MEC on PODS-loaded microwells expressed milk-associated proteins, we did not observe robust reconstitution of a bilayered epithelium. Higher-resolution printing at closer-to-physiological feature sizes was, on its own, insufficient to promote continuous bilayer formation. This suggests that lactogenic marker induction and epithelial stratification can be partially decoupled, and that bilayer formation likely requires additional factors beyond geometry, such as fluid flow or luminal-to-basal cell ratios.[46] Here, a key consideration for future work is donor-to-donor variability in the cellular composition and lactogenic responsiveness of milk MEC. In our previous work, flow-cytometric profiling of milk MEC has shown substantial inter-individual heterogeneity in EpCAM^+^ (luminal) and CD49f^+^ (basal) cells, which propagated during culture.[17] Therefore, an imbalanced starting population for scaffold seeding may prevent bilayer formation. Inter-donor variance could be addressed by directly controlling the input cell proportions. Donors with comparable baseline compositions and PRL responsiveness could be selected or cells could be sorted into luminal and basal populations and recombined at defined ratios to improve architectural fidelity. At the same time, this engineering objective should be interpreted in the context of native tissue organization. The literature indicates that basal coverage is relatively continuous around ducts but can be discontinuous around alveolar units, which can permit direct luminal contact with the basement membrane.[47,48] Thus, while controlling luminal:basal input ratios may be essential for reproducible architecture across donors, the appropriate endpoint for alveolar-like constructs may not be a perfectly continuous “textbook” bilayer. Instead, cell composition could be treated as a tunable design parameter, where luminal:basal ratios are systematically varied, and basal coverage is evaluated alongside lactogenic output to identify configurations that best support stable, long-term lactation-relevant epithelial constructs *in vitro*.

Together, these results position the platform as a tool to identify design rules for human mammary microtissues, where geometry, matrix composition, and endocrine delivery can be tuned independently in a standardized Transwell format. The epithelial-lined open lumen provides apical access, and together with the basolateral reservoir creates two independently addressable compartments that enable more complex functional assays, such as repeated sampling of apical products and barrier/transport measurements. Additionally, this compartmentalized set-up also provides a framework to interrogate of ductal-alveolar dysfunction, such as loss of polarity, barrier breakdown, or perturbed flow, processes implicated in lactational mastitis, an inflammatory disorder affecting up to one in three breastfeeding women worldwide.[49] Marked by pain, systemic symptoms, reduced milk production, and altered milk composition, mastitis highlights the need for more experimental models that recapitulate the 3D architecture of lactating alveoli *in vitro* and allow controlled stimulation or perturbation of the mammary microenvironment.[49] Looking ahead, improved z-axis control, higher curvature coverage, expansion to additional donors, and defined luminal:basal input ratios should enable a clearer understanding of how microarchitecture and transport shape epithelial organization and lactation-associated output. More broadly, our FLight scaffolds could be adapted to other secretory or barrier epithelia where polarity and directional readouts are central.

## 4. Experimental Section and Methods

### Bovine tissue collection, decellularization and ECM extraction

Fresh lactating bovine udders were processed and decellularized based on previously established protocols. [17] The tissue was cut into cubes ( [ 1.5 mm^3^) and washed in MiliQ water for 2h, then washed with a 1% SDS (Sigma, 75746) for 2h and finally decellularized in fresh 1% SDS (Sigma, 75746) for 16h (overnight) at room temperature, adding 2% Pen/Strep. The pieces were transferred to isopropanol for 5h (frequent changes), before washing in MiliQ water. The decellularized tissue was freeze-dried and cryomilled (Retsch) into a fine powder. The powder was enzymatically digested with 1 mg/ml pepsin (Sigma, P7012-250MG), in 0.1M HCL for 48h and then brought to neutral pH on ice. The neutralized dECM_mam_ was centrifuged at 8000 rpm for 7 min, freeze dried and stored at -20°C.

### Resin preparation

Col-I (Advanced Biomatrix, 5005) was resuspended at 10 mg/ml in phosphate- and glucose-rich buffer at 4°C. 40 mg/ml of the lyophilized dECM_mam_ were resuspended in 1X PBS on ice. Matrigel (Growth Factor Reduced, Corning, 356230) readily available from the provider at [ 10 mg/ml. 0.1mM RU and 10 mM SPS were added to the freshly prepared mixed resins (composed of 5 mg/ml Col-I, 5mg/ml Matrigel and/or 20 mg/ml dECM_mam_) in the dark on ice.

### Photorheology and stress relaxation

Rheology was performed using an Anton Paar MCR 302e rheometer with a 20 mm parallel plate geometry and a glass floor. The rheometer was combined with the Omnicure Series1000 lamp (Lumen Dynamics) with sequential 400–500 nm and narrow 405 nm bandpass filters (Thorlabs). Measurements were performed in the dark for 3 min before irradiating the sample with 405 nm light at 10 mW cm^−2^ intensity. Shear measurements were performed at a shear rate of 2% and a frequency of 1 Hz, with 10 seconds acquisition intervals. 76 μL of the sample was loaded onto the rheometer, with a gap distance of 0.2 mm and to prevent drying during testing, a wet tissue paper was added in the chamber.

### Mechanical testing

Cylindrical hydrogels were prepared using the FLight Top Down Projection into a 96 well plate (construct dimensions: 6 mm diameter; 2 mm height), washed and postcured in a UV box. The TA.XT Texture Analyzer (Stable Micro System) with a 500g load cell was used for compression tests. The cylindrical hydrogels were placed between compression plates, and a pre-load of 0.2 g was applied for full contact of the samples with the plates. Samples were compressed to 15% strain at 0.01mm/s. The compressive modulus was calculated by linear fitting the first 3% of the stress-strain curve.

### Dose tests

The printability of the mammary photoresins were assessed with the built-in software feature of the open format volumetric printer (Readily3D® SA, CH). Dose tests were performed by projecting [ 1 mm circles with increasing light intensity onto a 1 mm path length cuvette filled with the cooled photoresin mixes used in this study.

### Filamented Light (FLight) printing

The photoresins were pipetted into standard lab ware 48 well plates and 24 well Transwell inserts. FLight printing was perfomed with a custom built FLight printer. A fiber coupled laser (405 n, 1000 mW power) with a high speckle contrast ratio (for modulation instability enabling microfilament formation), was expanded with a 4 f magnification system and projected onto a digital micromirror array device (DMD; DLP6500FYE, Texas instruments) with a pixel resolution 1920 × 1080 and a pixel pitch 7.56 *μ*m.[50] The DMD allowed to project the light in digitally designed images (Microsoft PowerPoint for Microsoft 365), through a plano convex lenses (1:1 magnification) with an aperture in between (for isolation of the reflected image from the DMD diffraction pattern), onto a mirror for top-down Flight printing. A light intensity of 41.83 mW cm^−2^ was used. After printing, uncrosslinked photoresin was removed with cold PBS.

### Analysis of microstructures (filaments and microchannels)

Fluorescent z-stacks were acquired using an FVOlympus 3000 confocal microscope and images were analyzed in Fiji/ImageJ 1.54f (Java 1.8.0_332). For microfilament segmentation the default threshold was applied to all scans. The “straight line” function was used to draw a line perpendicular to the microfilaments/void spaces. The diameters of void spaces were measured using the “Plot Profile” plugin in Fiji software (n = 3, dataset size = 20). The “OrientationJ” plugin (available at http://bigwww.epfl.ch/demo/orientationj) was used to evaluate the filament orientation. [23] The structure tensor of the local window was set to 2 pixels and a “Gaussian” gradient was set.

### Human milk mammary epithelial cell isolation and expansion, and MCF10A culture

Human donors were recruited in line with Swiss Ethics guidelines and written informed consent was obtained from all participants (Kantonale Ethikkomission, 2022-02012). Human milk was collected and cells were isolated based on our previously established protocol. [17] Briefly, cells were isolated by centrifugation and the pellet was resuspended in MECGM (PromoCell, C-21010) with 10 uM Forskolin (HelloBio, HB1348), 3 uM Rock-I (Y-27632) (HelloBio, HB2297) and 1% Pen/Strep (Gibco,15140122). Cells were expanded on with Matrigel (Corning, 11573620)-coated polystyrene surfaces for 5 days before changing the medium to MECGM with 10 uM Forskolin, 0.5 % Human Serum (Sigma, H4522) and 1% Pen/Strep, at 37°C, at 5% O2. For further passaging cells were cultured in MECGM with 10 uM Forskolin, 5 % Human Serum and 1% Pen/Strep with media changes every 2 days until day 4 and daily after day 4 of culture. Cells were split at 80-90% confluency, with TryplE (Gibco, 12604021). MCF10A cells were cultured using established protocols. [36,p.10]

### Construct seeding and culture

20-40K cells were seeded onto the printed scaffolds and cultured at 37°C, at 5% O2, with daily media changes.

### PRL-PODS®

Custom slow release microcrystals were manufactured by Cell Guidance Systems Ltd. Recombinant human prolactin (PRL; UniProtKB accession P01236) was used for protein loading. The signal sequence was omitted and a tag at the N terminus was incorporated. The PRL amino-acid sequence was: MLPICPGGAA RCQVTLRDLF DRAVVLSHYI HNLSSEMFSE FDKRYTHGRG FITKAINSCH TSSLATPEDK EQAQQMNQKD FLSLIVSILR SWNEPLYHLV TEVRGMQEAP EAILSKAVEI EEQTKRLLEG MELIVSQVHP ETKENEIYPV WSGLPSLQMA DEESRLSAYY NLLHCLRRDS HKIDNYLKLL KCRIIHNNNC

### PODS® PRL release (ELISA)

To quantify PRL release, PODS®-laden FLight hydrogels were incubated in the mammary cell culture medium described above for 7 days. At days 1, 3, 5, and 7, 300 μL of culture medium was collected and stored at -20°C until used. PRL release was with a Human Prolactin Elisa Kit (Proteintech, KE00172) according to manufacturer’s protocols

### Prolactin structure retrieval and visualization

The human PRL NMR solution structure (PDB ID: 1N9D) was retrieved from the RCSB Protein Data Bank and downloaded as a PDB file on 24.07.25. Structures were visualized and figures prepared in PyMOL (The PyMOL Molecular Graphics System, Version 3.0 Schrödinger, LLC.); MODEL 1 was used for figure generation.[35]

### Rhodamine and RU/SPS diffusion

10 kDa Rhodamine B isothiocyanate-Dextran (R8881-100MG, Sigma) was dissolved in PBS to prepare a 10 mg/mL stock solution. 60 μL microliter of the stock were added to 1mL of the resins. Resins were printed the FLight printer at their respective light doses (625.5 mJ cm^−2^ for dECM_mam_+Matrigel, 334.6 mJ cm^−2^ for dECM_mam_+Col-I, and 125.5 mJ cm^−2^ for Matrigel+Col-I) and bulk controls were crosslinked in an UV box (at the same light doses, 405 nm). The printed hydrogel cylinders were placed in 1 mL of PBS. At designated time points, 500 μL of the supernatants were collected and replenished with fresh PBS. To quantify Rho release, fluorescence measurements were conducted at an excitation wavelength of 540 nm and an emission wavelength of 580 nm. For the release of RU/SPS the absorption was measured at 450 nm. The cumulative release over all time points was calculated.

### Immunohistochemistry and confocal imaging

Scaffolds with cells were fixed in 4% PFA for 1h, before immunofluorescent staining. The constructs were washed 3 times in PBS, and blocked with 5% bovine serum albumin (BSA, Millipore Sigma) with 0.2% Triton-X100 (Sigma, T8787-100ml) in PBS for 35 min at room temperature. The constructs were incubated with primary antibodies (**Table S2**, Supplementary Information) in BSA-PBS overnight at 4°. On the next day, samples were washed 3 times in PBS and incubated with the secondary antibody solution (**Table S3**, Supplementary Information) and DAPI diluted in BSA-PBS for 2h at room temperature. Samples were washed 3 times in PBS and imaged using a FVOlympus 3000 confocal microscope.

### Lipid droplet staining

Nile Red (Sigma, 19123) was resuspended in DMSO to prepare a 10 mM stock solution (3.18 mg/mL). For the staining solution the stock was diluted in PBS to 10 μM and Hoechst was added at a 1:1000 dilution. Constructs were washed with PBS for 10 minutes, incubated with the staining solution for 15 minutes, washed twice with PBS, and imaged using an Olympus FV4000 confocal microscope.

### Light sheet microscopy

An axially scanned light sheet microscope (MesoSPIM, V4) was used to image FLight printed constructs [51]. The constructs were transferred to a 4 mm glass cuvette with PBS that was inserted into a custom 3D-printed sample holder and submerged in a quartz chamber filled with miliQ water, mounted onto the MesoSPIM microscope stand. For imaging, a macro-zoom system (Olympus MVX-10) and 2x air objective (Olympus MVPLAPO1x) were used. Voltage adjustments using the electrically tunable lens (ETL) were performed. Step size 15 μm.

### RNA extraction and real-time qPCR

MCF10A for 5 days on FLight crosslinked disks (n=3 for each hydrogel formulation and TC plastic control). Cells were incubated for 10 min in BLT buffer and then mixed with 70% EtOH before transfer to the RNeasy mini kit (Qiagen, 74104) column to extract the total RNA. 100 ng of RNA was retrotranscribed with GoScript Reverse Transcriptase kit (Promega A5003, Madison, Wisconsin, USA). cDNA was diluted 1:5 with RNAse-free water. RT-qPCR was performed with GoTaq qPCR Master Mix (Promega, A6002, Madison, Wisconsin, USA). Reactions were run on a QuantStudio 3 96-well 0.1 ml Block Real-Time PCR System (Applied Biosystems™, Waltham, Massachusetts, USA). Samples were analyzed using amplification and melting curves. Ct values were normalized to GAPDH and to the TC plastic controls. All RT-qPCR primer pairs designed for this study were bought through Microsynth AG (Balgach, Switzerland) and can be found in **Table S4** (Supplementary Information).

### Western Blotting

Milk MEC were grown on 3D printed constructs and collected with TryplE on days 5 or 7. Cells were lysed in RIPA buffer with protease inhibitors (Sigma-Aldrich P1860-1ML, Burlington, Massachusetts, USA) and then centrifuged for 10 min at 12,000□×□g. The resulting proteins were mixed with NuPAGE™ Sample Reducing Agent (Invitrogen™ NP0004, Waltham, Massachusetts, USA) and NuPAGE™ LDS Sample Buffer (Invitrogen™ NP0007, Waltham, Massachusetts, USA) and denatured for 10 min at 80°C in a thermocycler. Samples were run on a NuPAGE™ 4–12%, Bis–Tris, 1.0–1.5 mm, Mini Protein Gel (Thermo Scientific™ NP0321BOX, Waltham, Massachusetts, USA) and transferred onto a nitrocellulose membrane. The membrane was incubated overnight (at 4°C) with primary antibodies (**Table S5**, Supplementary Information). Membranes were washed with PBS supplemented with 0.1% Tween and incubated with a secondary HRP-conjugated goat anti-rabbit or anti-mouse IgG antibodies. The HRP signal was detected with WesternBright ECL HRP substrate (Advansta K-12045-D20, San Jose, California, USA) and imaged with a FUSION FX6 EDGE Imaging System (Witec, Sursee, Switzerland).

### Quantification and statistical analysis

The sample size (n) for each experiment is provided in the corresponding figure legends. Data analysis was performed using Excel (Microsoft 365 MSO Version 2412 Build 16.0.18324.20092 64-bit), Matlab R2018a (9.4.0.813654), GraphPad Prism (version 10.4.0 for Windows) and Fiji/ImageJ 1.54f (Java 1.8.0_332). Data are presented as mean ± SD, and statistical significance was assessed using a paired t-test and p-value < 0.05 was considered statistically significant. Only significant results are shown in the figure panels. Object circularity: was quantified in Fiji/ImageJ (v1.54f) using the built-in shape descriptors. Following image segmentation, regions of interest were measured using Analyze > Measure and/or Analyze > Analyze Particles with “Circularity” enabled in Analyze > Set Measurements. Circularity was computed as 4nA/P^2^; values range from 1.0 for an ideal circle toward 0 for increasingly elongated morphologies.

### Graphical abstract and proofreading tools

The graphical abstract consists solely of schematic illustrations and does not present original experimental data or results. Schematics for the graphical abstract were created using “Adobe Fresco: Draw & Paint” App (Adobe Inc., Version 7.1.0) on an iPad (Apple, 10th generation), PowerPoint (for Microsoft 365) and Biorender (https://BioRender.com). A large language model (ChatGPT, OpenAI) was used to generate a draft graphical element (well-plate icon) and to assist with proofreading; all text and figures were verified and finalized by the authors.

## Acknowledgments

We greatly appreciate and thank the participating mothers for their milk sample donations. We thank Simon Moser, Shannon Kelleher, Parth Chansoria, Hao Liu and all the members of the laboratory for the discussions and inputs. We are grateful to Jakub Janiak for creating the 3D rendering of the FLight printer and building it for our laboratory. We thank Kristian Ivkovic for kindly providing feedback on the manuscript. The authors gratefully acknowledge ScopeM for their support and assistance in this work. Lightsheet imaging was performed with equipment maintained by the Center for Microscopy and Image Analysis, University of Zurich, with special thanks to Nikita Vladimirov and José María Mateos.

## Data and materials availability

All data needed to evaluate the conclusions in the paper are present in the paper and/or the Supplementary Materials. Data also is available in the ETH Zurich Research Collection (https://doi.org/10.3929/ethz-c-000794688) under the terms of the repository’s data-sharing policies.

## References

[1] Truchet S, Honvo-Houéto E. Physiology of milk secretion. Best Practice & Research Clinical Endocrinology & Metabolism. 2017;31(4):367–384.

[2] Slepicka PF, Somasundara AVH, dos Santos CO. The molecular basis of mammary gland development and epithelial differentiation. Seminars in Cell & Developmental Biology. 2021;114:93–112.

[3] Biswas SK, Banerjee S, Baker GW, et al. The Mammary Gland: Basic Structure and Molecular Signaling during Development. International Journal of Molecular Sciences. 2022;23(7):3883.

[4] Adriance MC, Inman JL, Petersen OW, et al. Myoepithelial cells: good fences make good neighbors. Breast Cancer Res. 2005;7(5):190.

[5] Shennan DB, Peaker M. Transport of Milk Constituents by the Mammary Gland. Physiological Reviews. 2000;80(3):925–951.

[6] Neville MC, McFadden TB, Forsyth I. Hormonal Regulation of Mammary Differentiation and Milk Secretion. Journal of Mammary Gland Biology and Neoplasia. 2002;

[7] Rooney N, Streuli CH. How integrins control mammary epithelial differentiation: A possible role for the ILK–PINCH–Parvin complex. FEBS Letters. 2011;585(11):1663–1672.

[8] LeMaster C, Pierce SH, Geanes ES, et al. The cellular and immunological dynamics of early and transitional human milk. Communications Biology. 2023;6:539.

[9] Muschler J, Lochter A, Roskelley CD, et al. Division of Labor among the α6β4 Integrin, β1 Integrins, and an E3 Laminin Receptor to Signal Morphogenesis and β-Casein Expression in Mammary Epithelial Cells. Molecular Biology of the Cell. 1999;10(9):2817.

[10] Nelson CM, Bissell MJ. Modeling dynamic reciprocity: Engineering three-dimensional culture models of breast architecture, function, and neoplastic transformation. Seminars in Cancer Biology. 2005;15(5):342–352.

[11] Alcaraz J, Xu R, Mori H, et al. Laminin and biomimetic extracellular elasticity enhance functional differentiation in mammary epithelia. EMBO J. 2008;27(21):2829–2838.

[12] Rosenbluth JM, Schackmann RCJ, Gray GK, et al. Organoid cultures from normal and cancer-prone human breast tissues preserve complex epithelial lineages. Nat Commun. 2020;11(1):1711.

[13] Bissell MJ, Rizki A, Mian IS. Tissue architecture: the ultimate regulator of breast epithelial function. Curr Opin Cell Biol. 2003;15(6):753–762.

[14] Lu L, Fuji K, Guyomar T, et al. Generic comparison of lumen nucleation and fusion in epithelial organoids with and without hydrostatic pressure. Nature Communications. 2025;16:6307.

[15] Kim D, Lim H, Youn J, et al. Scalable production of uniform and mature organoids in a 3D geometrically-engineered permeable membrane. Nat Commun. 2024;15(1):9420.

[16] Gjorevski N, Nikolaev M, Brown TE, et al. Tissue geometry drives deterministic organoid patterning. Science. 375(6576):eaaw9021.

[17] Hasenauer A, Bevc K, McCabe MC, et al. Volumetric printed biomimetic scaffolds support in vitro lactation of human milk-derived mammary epithelial cells. Science Advances [Internet]. 2025 [cited 2025 Sep 29]; doi: 10.1126/sciadv.adu5793.

[18] Chanson L, Brownfield D, Garbe JC, et al. Self-organization is a dynamic and lineage-intrinsic property of mammary epithelial cells. Proceedings of the National Academy of Sciences. 2011;108(8):3264–3269.

[19] Luciano M, Versaevel M, Kalukula Y, et al. Mechanoresponse of Curved Epithelial Monolayers Lining Bowl-Shaped 3D Microwells. Advanced Healthcare Materials. 2024;13(4):2203377.

[20] Kim S-H, Lee GH, Park JY. Microwell fabrication methods and applications for cellular studies. Biomed Eng Lett. 2013;3(3):131–137.

[21] Guo W, Chen Z, Feng Z, et al. Fabrication of Concave Microwells and Their Applications in Micro-Tissue Engineering: A Review. Micromachines. 2022;13(9):1555.

[22] Rizzo R, Ruetsche D, Liu H, et al. Optimized Photoclick (Bio)Resins for Fast Volumetric Bioprinting. Advanced Materials. 2021;33(49):2102900.

[23] Liu H, Chansoria P, Delrot P, et al. Filamented Light (FLight) Biofabrication of Highly Aligned Tissue-Engineered Constructs. Advanced Materials. 2022;34(45):2204301.

[24] Liu H, Scherpe L, Hummer LB, et al. Filamented Light (FLight) Biofabrication of Mini-Tendon Models Show Tunable Matrix Confinement and Nuclear Morphology [Internet]. bioRxiv; 2025 [cited 2025 Feb 26]. p. 2025.02.03.636123. Available from: https://www.biorxiv.org/content/10.1101/2025.02.03.636123v1.

[25] Puiggalí-Jou A, Rizzo R, Bonato A, et al. FLight Biofabrication Supports Maturation of Articular Cartilage with Anisotropic Properties. Adv Healthc Mater. 2024;13(12):e2302179.

[26] Winkelbauer M, Hasenauer A, Rütsche D, et al. Rapid Deep Vat Printing Using Photoclickable Collagen-Based Bioresins. Advanced Healthcare Materials. 2025;14(28):2405105.

[27] Pelham RJ, Wang Y. Cell locomotion and focal adhesions are regulated by substrate[flexibility. Proceedings of the National Academy of Sciences. 1997;94(25):13661–13665.

[28] Hughes K. Comparative mammary gland postnatal development and tumourigenesis in the sheep, cow, cat and rabbit: Exploring the menagerie. Seminars in Cell & Developmental Biology. 2021;114:186–195.

[29] Twigger A-J, Engelbrecht LK, Bach K, et al. Transcriptional changes in the mammary gland during lactation revealed by single cell sequencing of cells from human milk. Nat Commun. 2022;13(1):562.

[30] Martin Carli JF, Trahan GD, Jones KL, et al. Single cell RNA sequencing of human milk-derived cells reveals sub-populations of mammary epithelial cells with molecular signatures of progenitor and mature states: a novel, non-invasive framework for investigating human lactation physiology. J Mammary Gland Biol Neoplasia. 2020;25(4):367–387.

[31] Ravasio A, Cheddadi I, Chen T, et al. Gap geometry dictates epithelial closure efficiency. Nat Commun. 2015;6(1):7683.

[32] Zhao B, Li L, Lu Q, et al. Angiomotin is a novel Hippo pathway component that inhibits YAP oncoprotein. Genes Dev. 2011;25(1):51–63.

[33] Liu X, Robinson GW, Gouilleux F, et al. Cloning and expression of Stat5 and an additional homologue (Stat5b) involved in prolactin signal transduction in mouse mammary tissue. Proceedings of the National Academy of Sciences. 1995;92(19):8831–8835.

[34] Bridgewater RE, Streuli CH, Caswell PT. Extracellular matrix promotes clathrin-dependent endocytosis of prolactin and STAT5 activation in differentiating mammary epithelial cells. Sci Rep. 2017;7(1):4572.

[35] Keeler C, Dannies PS, Hodsdon ME. The Tertiary Structure and Backbone Dynamics of Human Prolactin. Journal of Molecular Biology. 2003;328(5):1105–1121.

[36] Qu Y, Han B, Yu Y, et al. Evaluation of MCF10A as a Reliable Model for Normal Human Mammary Epithelial Cells. PLoS ONE. 2015;10(7):e0131285.

[37] Rios AC, Fu NY, Jamieson PR, et al. Essential role for a novel population of binucleated mammary epithelial cells in lactation. Nature Communications. 2016;7:11400.

[38] Vasquez CG, Vachharajani VT, Garzon-Coral C, et al. Physical basis for the determination of lumen shape in a simple epithelium. Nat Commun. 2021;12(1):5608.

[39] Bovyn MJ, Haas PA. Shaping epithelial lumina under pressure. Biochemical Society Transactions. 2024;52(1):331–342.

[40] Cerruti B, Puliafito A, Shewan AM, et al. Polarity, cell division, and out-of-equilibrium dynamics control the growth of epithelial structures. Journal of Cell Biology. 2013;203(2):359–372.

[41] Ferrell N, Desai RR, Fleischman AJ, et al. A microfluidic bioreactor with integrated transepithelial electrical resistance (TEER) measurement electrodes for evaluation of renal epithelial cells. Biotechnology and Bioengineering. 2010;107(4):707–716.

[42] Guan X, Sun W, Ma Z, et al. From lab to industry: Technologies and challenges for scaling up bioprocesses in cell-based food production. Trends in Food Science & Technology. 2025;161:105040.

[43] Jochems PGM, van Bergenhenegouwen J, van Genderen AM, et al. Development and validation of bioengineered intestinal tubules for translational research aimed at safety and efficacy testing of drugs and nutrients. Toxicology in Vitro. 2019;60:1–11.

[44] Rizzo R, Rütsche D, Liu H, et al. Multiscale Hybrid Fabrication: Volumetric Printing Meets Two-Photon Ablation. Advanced Materials Technologies. 2023;8(11):2201871.

[45] Lee G, Lee J, Oh H, et al. Reproducible Construction of Surface Tension-Mediated Honeycomb Concave Microwell Arrays for Engineering of 3D Microtissues with Minimal Cell Loss. PLOS ONE. 2016;11(8):e0161026.

[46] Buchholz M-B, Bernal PN, Bessler N, et al. Development of a bioreactor and volumetric bioprinting protocol to enable perfused culture of biofabricated human epithelial mammary ducts and endothelial constructs. Biofabrication. 2025;17(3):031001.

[47] Oakes SR, Hilton HN, Ormandy CJ. Key stages in mammary gland development - The alveolar switch: coordinating the proliferative cues and cell fate decisions that drive the formation of lobuloalveoli from ductal epithelium. Breast Cancer Research. 2006;8(2):207.

[48] Srivastava V, Hu JL, Garbe JC, et al. Configurational entropy is an intrinsic driver of tissue structural heterogeneity. bioRxiv. 2023;2023.07.01.546933.

[49] Wilson E, Woodd SL, Benova L. Incidence of and Risk Factors for Lactational Mastitis: A Systematic Review. Journal of Human Lactation. 2020;36(4):673.

[50] Puiggalí-Jou A, Hui I, Baldi L, et al. Biofabrication of anisotropic articular cartilage based on decellularized extracellular matrix. Biofabrication. 2025;17(1):015044.

[51] Voigt FF, Kirschenbaum D, Platonova E, et al. The mesoSPIM initiative: open-source light-sheet microscopes for imaging cleared tissue. Nat Methods. 2019;16(11):1105–1108.

