## Supplementary Material for "A 3D printed model of human lactation"

**ORCIDs:**

Amelia Hasenauer (<https://orcid.org/0000-0003-4512-6195>)

Marcy Zenobi-Wong ([https://orcid.org/0000-0002-8522-9909](https://orcid.org/0000-0002-6107-6848))

**This file includes:**

Figures S1 to S13

Tables S1 to S5

Any supplementary methods have been provided in the main manuscript.

**Table S1.** List of milk samples used in this study.

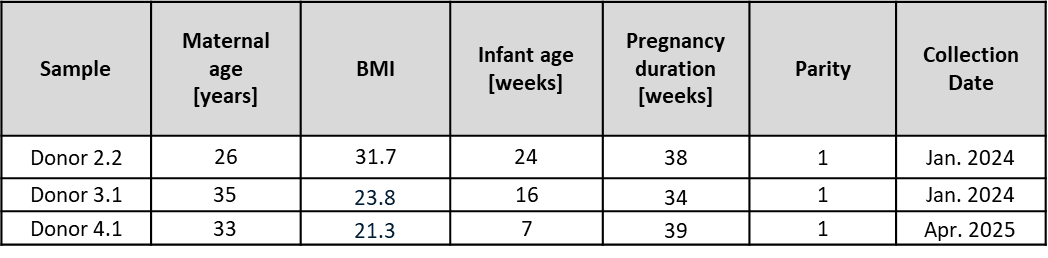

**Table S2. List of primary antibodies used for immunostaining**
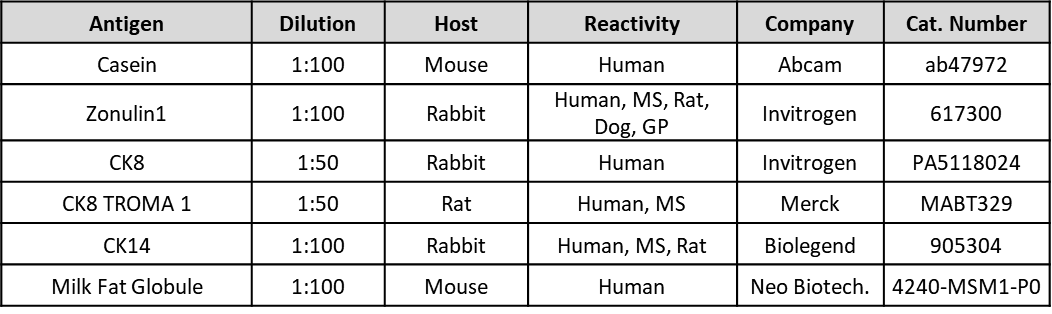

**Table S3. List of secondary antibodies used for immunostaining**
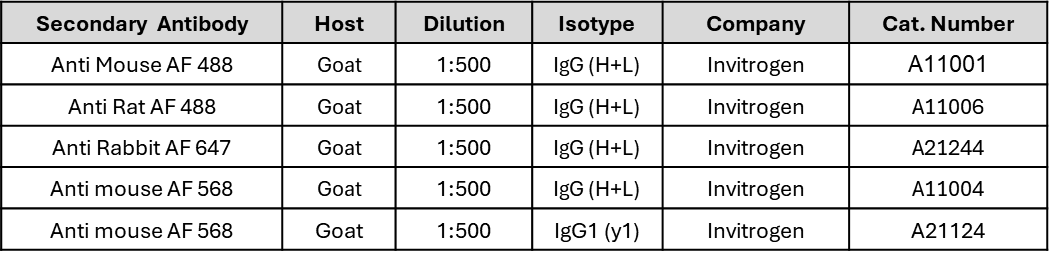

**Table S4. List of RT-qPCR primers**
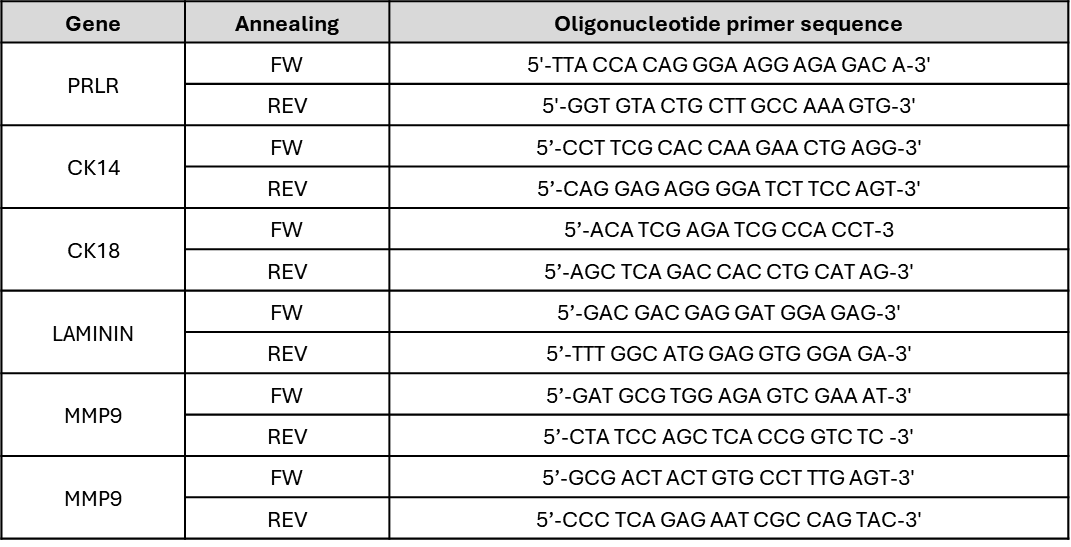

**Table S5. List of antibodies used for Western Blot**
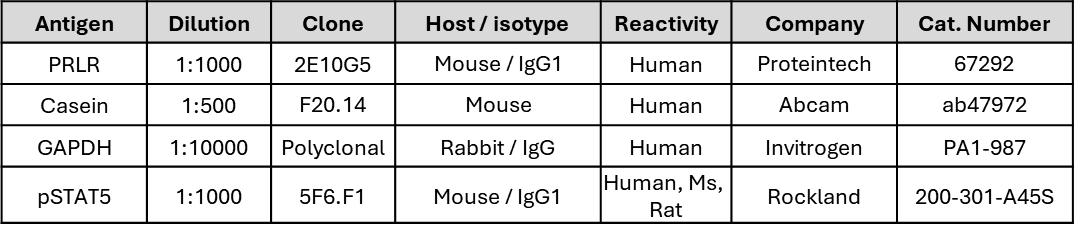

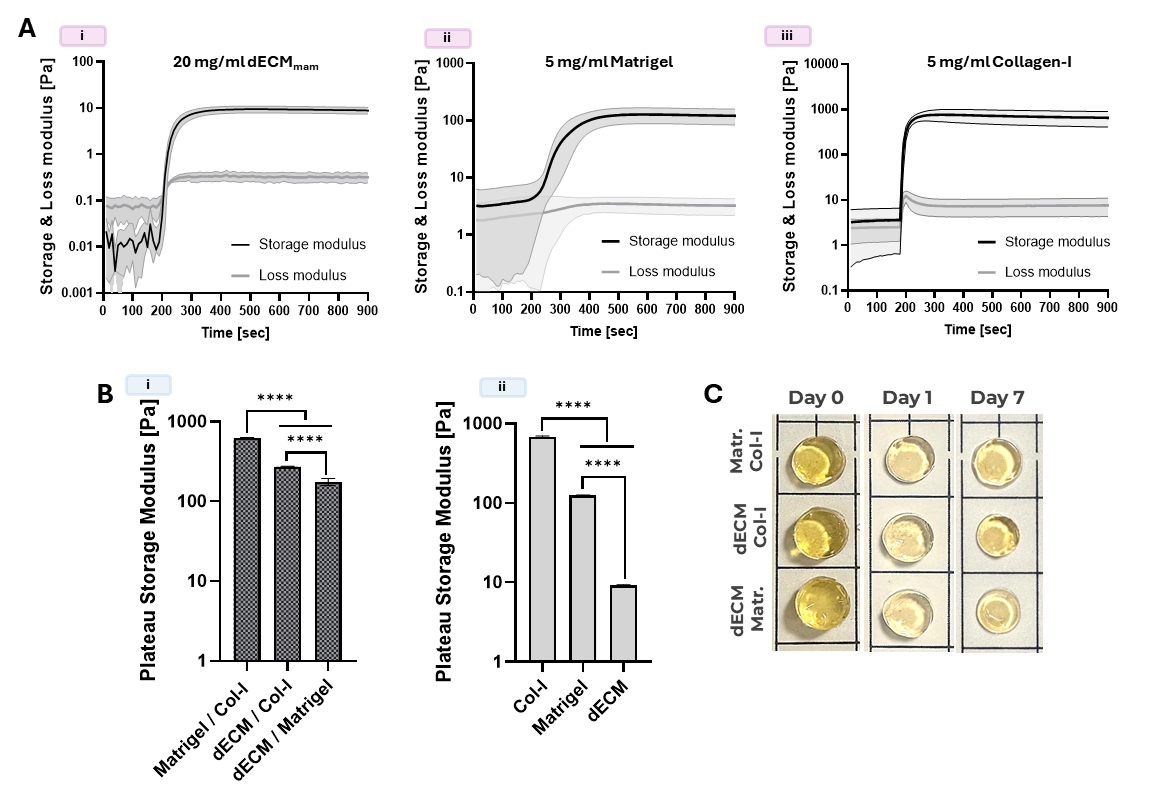

**Figure S1: Characterization of mammary ECM-based photoresins used for FLight printing.**

**A)** Photoreology showing time-dependent gelation during Ru/SPS-initiated photocrosslinking of the individual matrix components: **(i)** 20 mg/mL dECM_mam_, **(ii)** 5 mg/mL Matrigel, and **(iii)** 5 mg/mL collagen I. Storage modulus (G′, black) and loss modulus (G″, brey) are plotted over time. **B)** Plateau storage modulus (G′) of **(i)** the three ECM composite resins used in this study (dECM+Col-I, dECM+Matrigel) and **(ii)** the corresponding single-component resins (Col-I, Matrigel, dECM_mam_), all formulated with Ru/SPS (0.1 mM/10 mM). Mean ± S.D, n = 3, **** p<0.0001). **C)** Photographs of FLight printed cylindrical constructs at days 0, 1, and 7 in PBS at 37 °C (incubator), used to assess construct shrinkage/stability over time.

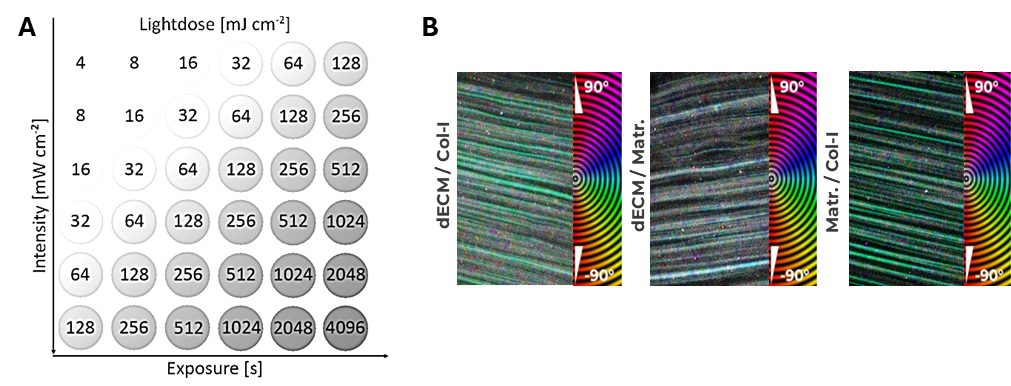

**Figure S2: Dose determination and filament orientation mapping.**

**A)** Light-dose matrix used to identify suitable filamented light (FLight) printing conditions by varying projection intensity (mW cm⁻²) and exposure time (s); each condition is labeled with the resulting delivered dose (mJ cm⁻²). **B)** Representative top-down images of FLight patterns in the indicated hydrogel formulations with corresponding orientation maps. Color encodes the local filament orientation angle (−90° to +90°) relative to the reference axis, as indicated by the color wheel.

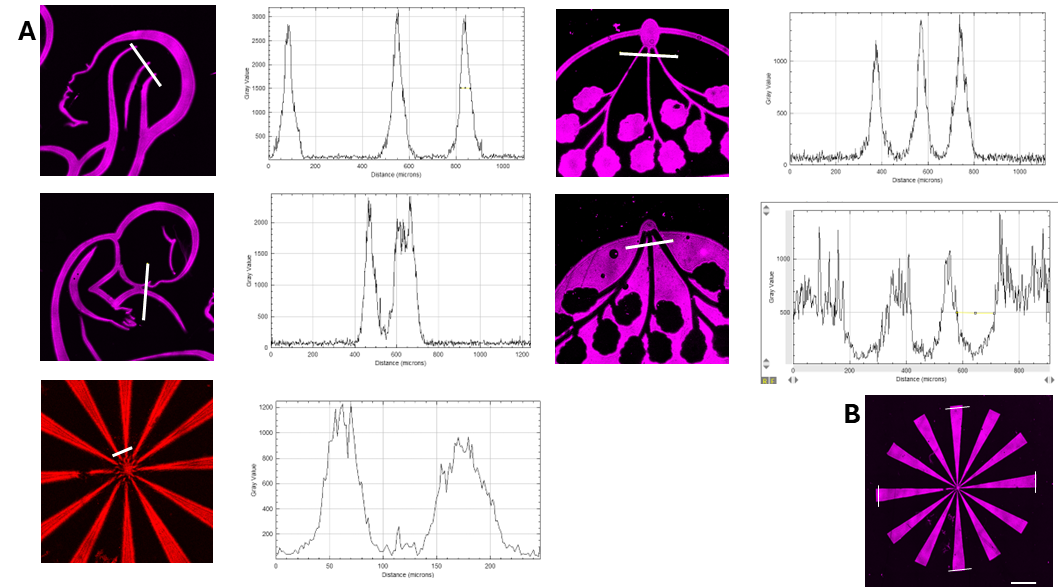

**Figure S3: Quantification of printing resolution and conformity.**

**A)** Printing resolution was quantified from 3D surface height maps by generating surface plots and line profiles across representative features (white line). Negative (recessed) resolution was defined as peak-to-peak valley spacing, and positive (raised) resolution as the width of a single peak. Representative Collagen–Matrigel examples are shown; the same analysis was applied to the other resins, and measurements were pooled across structures to obtain overall resolution distributions. **B)** Printing conformity was evaluated using the printed spoked wheel by measuring peripheral segment widths and comparing them to the nominal 500 µm design.

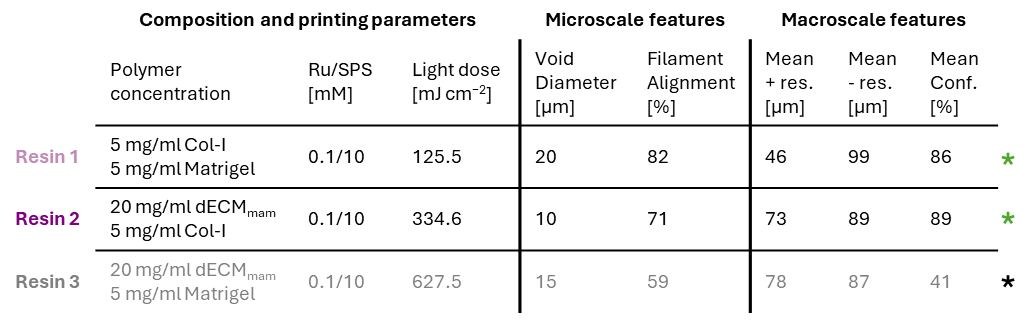

**Figure S4: Summary of the formulation and printing performance of three mammary hydrogel resin compositions in FLight printing.**

Three ECM-composite resins were compared: Resin 1 (Matrigel/Col-I), Resin 2 (dECM_mam_/Col-I), and Resin 3 (dECM_mam_/Matrigel), all formulated with Ru/SPS (0.1/10 mM). Left: composition and formulation-specific printing light dose (125.5, 334.6, and 627.5 mJ cm^-2^ for Resins 1–3, respectively). Middle: filament-scale microarchitecture (void diameter and filament alignment). Right: macroscale printing metrics (positive/negative feature resolution and overall conformity to the design). Based on higher print fidelity/conformity, Resins 1 and 2 (green asterisks) were selected for subsequent mammary tissue model experiments.

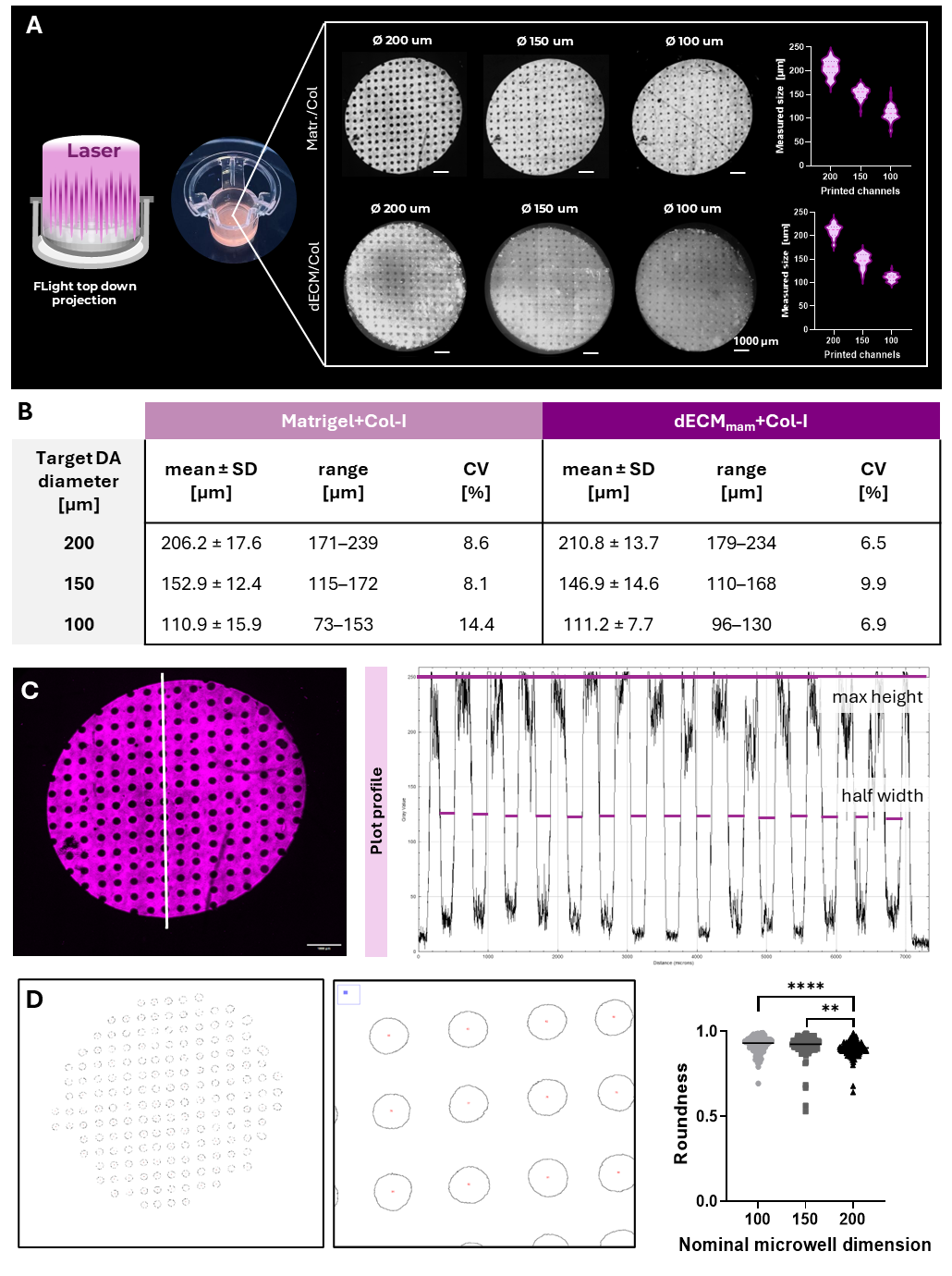

**Figure S5: Feature-size fidelity across designed microwell arrays.**

**A)** Representative top-view images of printed microwell/channel arrays fabricated in Matrigel/Col-I (top row) or dECM_mam_/Col-I (bottom row) with nominal target diameters (Ø= 200, 150, or 100 µm). Violin plots show the distributions of measured printed diameters for each design within each material. **B)** Summary of printed feature sizes for each target diameter and material, reported as mean ± SD, range, and coefficient of variation (CV = (SD/mean) × 100%.). **C)** Example measurement workflow used to extract feature diameters from top-view images: a line region-of-interest (ROI) was drawn across the microwell array (left), and the corresponding grayscale intensity plot profile (right) was used to determine feature width via a half-maximum (half-width) criterion relative to the local maximum intensity. **D)** Well “roundness” assessed in Fiji/ImageJ segmentation across printed microwell diameters in Matrigel/Col-I (Ø200 μm).

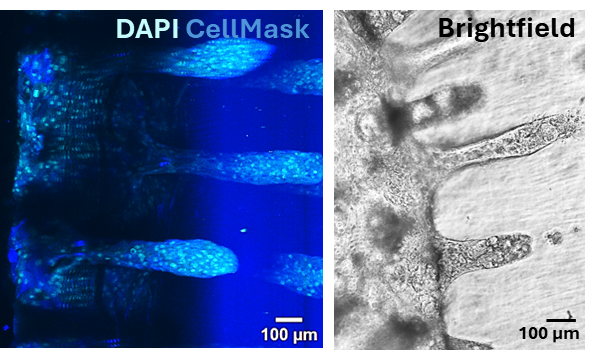

**Figure S6: Milk MEC form duct-like epithelial linings in deeper FLight-printed microwells.**

Milk-derived mammary epithelial cells (milk MEC) seeded on 1 mm–deep FLight-printed microwell scaffolds, where the increased resin fill height produces deeper, channel-like geometries that resemble ductal structures. **Left**: fluorescence image showing nuclei (DAPI, cyan) and CellMask-labeled cell membranes (blue) outlining epithelial coverage along the microwell walls. **Right**: representative brightfield image.

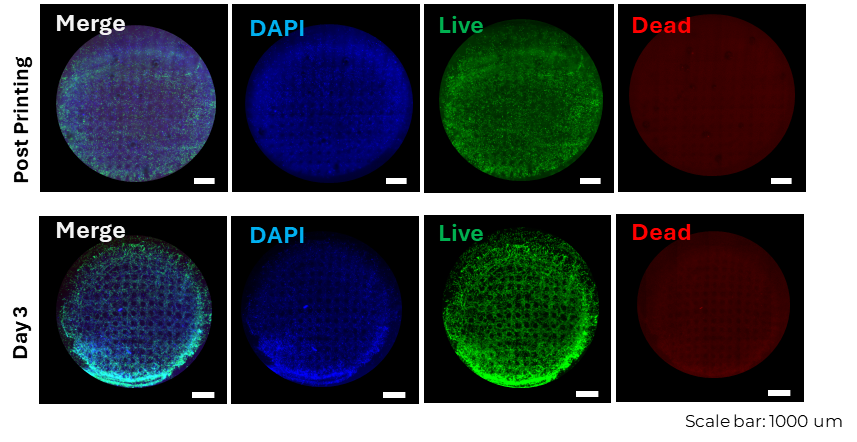

**Figure S7: Cytocompatibility of stromal-cell encapsulation in dECM_mam_/Collagen microwell scaffolds.**

Representative Live/Dead staining of 3T3 fibroblasts encapsulated within dECM_mam_/Col-I microwell scaffolds (nominal Ø200 µm wells) immediately after printing and after 3 days in culture. Shown are merged images and individual channels for nuclei (DAPI, blue), live cells (calcein-AM, green), and dead cells (propidium iodine, red).

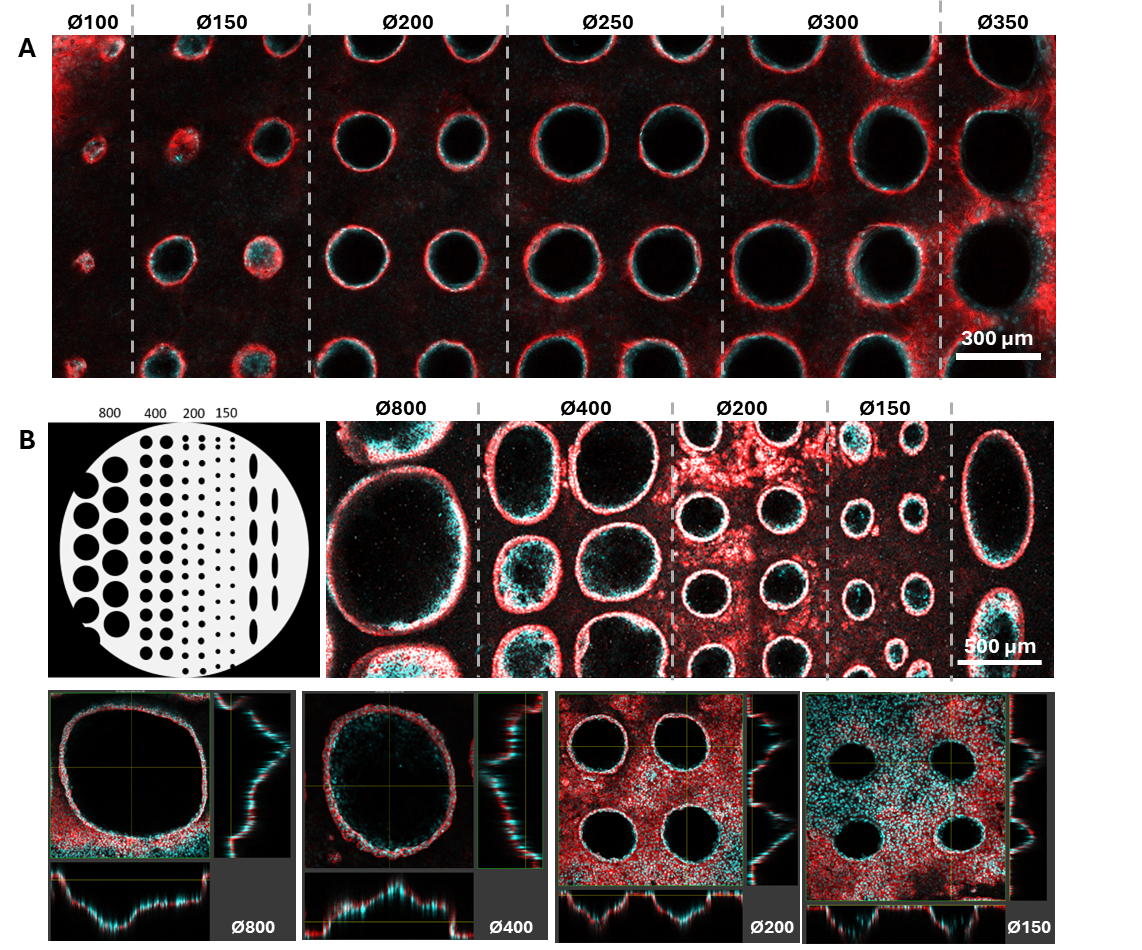

**Figure S8: Size-dependent lumen formation of milk MEC in multi-diameter microwell arrays.**

**A)** Fluorescence image of a FLight-printed insert containing adjacent regions of increasing microwell diameters (Ø100, Ø150, Ø200, Ø250, Ø300, Ø350 µm). Dashed lines mark the boundaries between diameter zones. Milk MEC preferentially line larger wells, whereas smaller wells more frequently show partial filling/occlusion **B)** Left: schematic of a multi-diameter design used for broader size screening (Ø800, Ø400, Ø200, Ø150 µm). Right: representative fluorescence image of milk MEC distributed across the diameter zones (dashed lines indicate zone boundaries). Bottom: orthogonal views for selected diameters (Ø800, Ø400, Ø200, Ø150 µm), illustrating size-dependent differences in epithelial coverage along the well walls and lumen openness. (Milk MEC were stained with DAPI (cyan, nuclei) and F-actin (red).

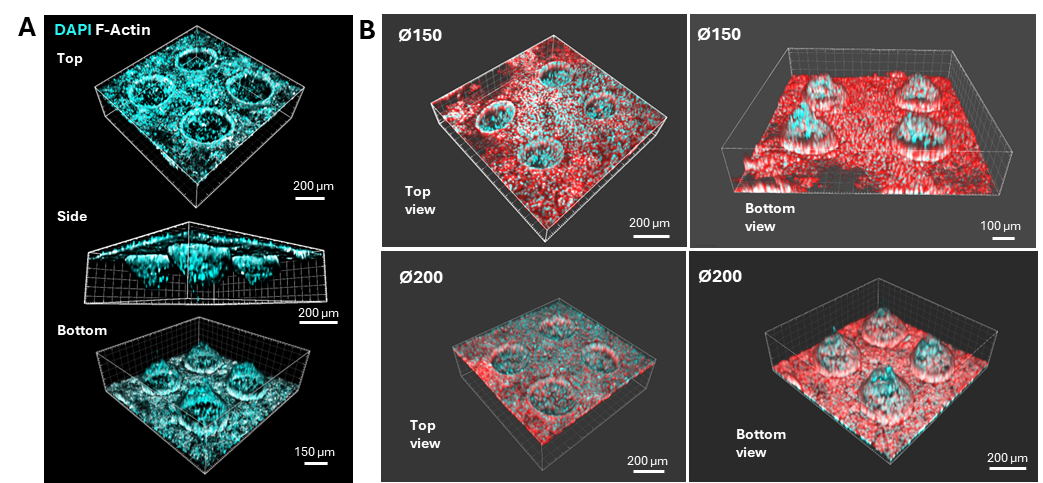

**Figure S9: Additional 3D imaging of milk MECs microwell arrays.**

Confocal z-stacks were reconstructed as 3D volume renderings to visualize cell coverage within the printed microwell geometry. Left: Renderings (top, side, and bottom views) of DA Ø 200. Right: Additional renderings of DA Ø150 and Ø200 microwell arrays viewed from the top and bottom, with F-actin (red) and nuclei (DAPI, cyan) to resolve cytoskeletal organization and nuclear positioning across the curved features.

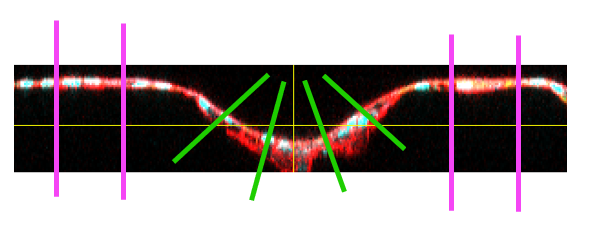

**Figure S10: Epithelial layer thickness in flat versus curved regions of printed microwells.** Representative confocal orthogonal section of a microwell seeded with milk MECs and stained for nuclei (DAPI, cyan) and F-actin (phalloidin, red). Green line segments indicate the line-scan positions used to quantify curved regions, whereas purple line segments indicate the positions used to quantify flat regions. For each microwell, measurements were performed four times per region by extracting fluorescence intensity line profiles and determining layer thickness as the full width at half maximum of the actin intensity peak.

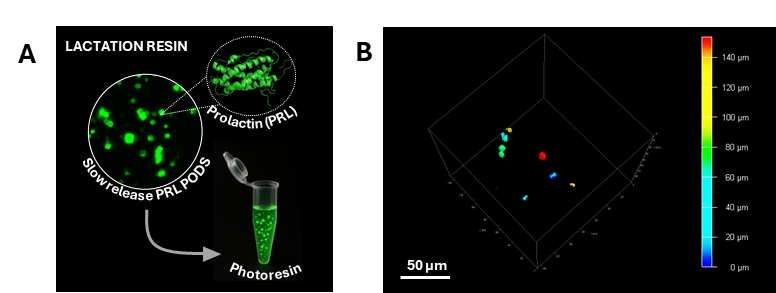

**Figure S11: Lactation resin formulation and PODS distribution.**

**A)** PRL PODS were mixed into the photoresin to generate the lactation resin; PRL NMR structure visualized in PyMol, representative fluorescence image to show dispersed PRL PODS (green) in resin, and schematic of an Eppendorf tube filled with lactation resin. **B)** After printing, a 3D z-stack reconstruction confirms PODS dispersion throughout the full construct depth; color denotes height (z, µm). Scale bar, 50 µm.

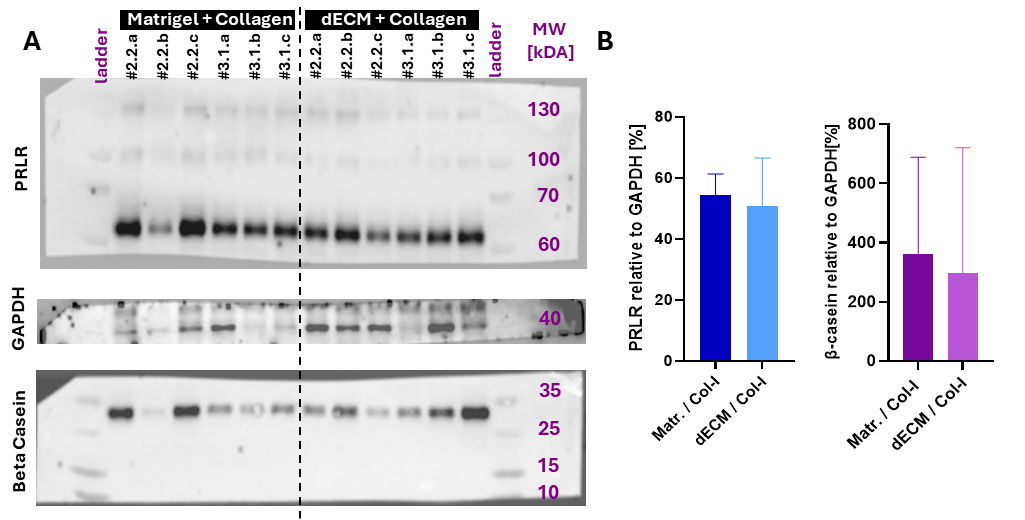

**Figure S12: Protein expression in Matrigel/Col-I versus dECM_mam_/Col-I conditions.** Representative Western blots of samples prepared from constructs generated using Matrigel/Col-I or dECM_mam_/Col-I formulations (left), with molecular weight markers (kDa) indicated. The dashed line denotes a splice separating lane groups from the two conditions. Lane labels correspond to the donors and the technical replicates (derived from an independent cell culture) are denoted as a, b, c,. GAPDH (~35–40 kDa) is shown as the loading control. Right: densitometric quantification of the indicated bands, normalized to GAPDH and plotted as mean ± SD.

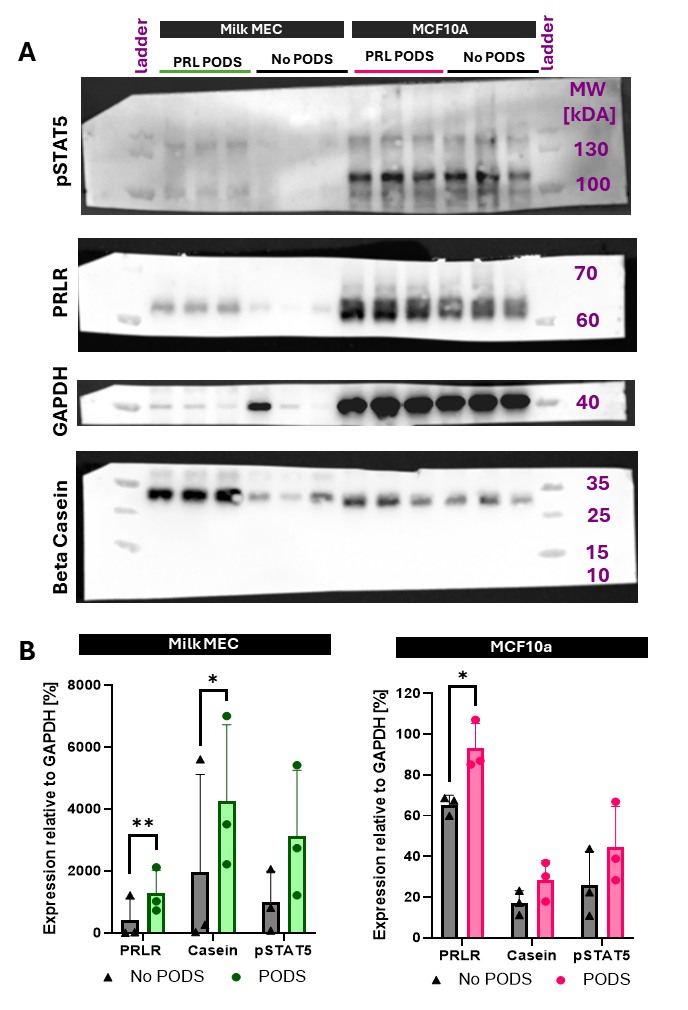

**Figure S13. Western blot analysis of Milk MEC (Donor 2.2) and MCF10A cells cultured in the Flight microwell system ± PODS.**

A) Western blots of MilkMEC and MCF10A lysates probed for PRLR, casein, and pSTAT5; molecular weight markers are indicated in purple. GAPDH served as the loading control. B) Quantification of PRLR, casein, and pSTAT5 normalized to GAPDH for each sample. Mean ± S.D, n = 3 independent cultures of Donor 2.2 and MCF10A cells, p<0.05*, p<0.01**
